# Correlations in population dynamics in multi-component networks

**DOI:** 10.1101/839019

**Authors:** Sonica Saraf, Lai-Sang Young

## Abstract

This is a theoretical study of correlations in spiking activity between neuronal populations. We focus on the spike firing of entire local populations without regard to the identities of the neurons that fire the spikes, and show that such a population-level metric is more robust than correlations between pairs of neurons. Between any source and target populations, there is an intrinsic response time characterized by the phase-shift that maximizes the correlation between their spiking. We find that the alignment of gamma-band rhythms contributes significantly to the positive correlations between populations. Hence, the correlation metric sheds light on the transference of gamma rhythms between populations; the effectiveness of such transference has been hypothesized to be connected to communication between brain regions. We investigate the dependence of correlations on connectivity and degree of synchrony, and consider multi-component network motifs with configurations known to occur in real cortex, studying the correlations between components that are directly or indirectly connected, by single or multiple pathways, with or without feedback. Mechanistic explanations are offered for many of the phenomena observed.

## Introduction

This paper reports on results of a theoretical study of correlations in spiking activity between local populations in the cerebral cortex. When a signal is transmitted from one region of the brain to another, one expects activity levels in the source and target regions to covary. Correlation is a measure of the effectiveness of this information transfer. As even fairly simple tasks involve multiple brain regions and signals are passed around in complicated ways, the relevance of correlations in activity level between brain regions requires little justification.

In this paper, we carry out a study using semi-realistic models. Our local populations are groups of a few hundred Excitatory and Inhibitory integrate-and-fire neurons, with connectivities similar to those in local circuits of the cerebral cortex. Population activity is driven by external input together with dynamical interaction among the neurons. We wanted our models to be realistic enough so that our results can be guided by experiments. At the same time, we do not wish to customize the study to specific brain regions, preferring to produce broadly relevant results. Three distinguishing features of our study are:

i. We focus on correlations in *spike firing* rather than on currents, because spike counts are a simple and direct measure of a region’s synaptic output.
ii. We study correlations in *population activity* without distinguishing between the spikes fired by individual neurons in the local population. This is a medium-size, low-variance statistic, one that we believe is functionally more relevant than correlations between specified pairs of neurons.
iii. We build *multi-component networks* following canonical motifs common in the cortex and study correlations between components that are directly as well as indirectly connected, with or without feedback, and with single or multiple pathways of connections.

A number of experimental correlation studies have been reported in the literature. This includes correlations between LFP in different regions of cortex (Jia et al. 2013; Salazar et al. 2012; Bosman et al. 2012; Jia et al. 2011), and between LFP and spike firing, as in spike-field correlations (Gray et al. 1989; Gray and Singer 1989; Zandvakili and Kohn 2015; Jia et al. 2013). Correlations in spiking behaviors between pairs of neurons have also been measured experimentally and are found to be very small (Zandvakili and Kohn 2015). Techniques such as “shuffling” and “jittering” have been devised to make these statistics more robust (Zandvakili and Kohn 2015; Jia et al. 2013). We propose that correlations in spiking on the population level are a natural alternative. We will connect these quantities, which are not easy to measure in the laboratory at the present time, to more accessible quantities by studying the dependence of correlations on sample size. One of the aims of this theoretical study is to inform and complement experimental measurements.

We have chosen to study correlated behaviors on relatively short timescales. As local-in-time dynamics are dominated by gamma-band activity, our study will in particular address the extent to which gamma patterns are transferred from one region to another. Some studies have suggested the importance of gamma power and coherence in relation to memory (Sederberg et al. 2003), perception, and sensory-motor tasks (Buzsáki 2011). Others have suggested that gamma rhythms are a by-product of neuronal processes, hence indicative of cortical state (Chariker et al. 2018). Both views support the relevance of coherence of gamma rhythms between brain regions. Because gamma rhythms are a result of population dynamics and are not reflected in the activity of individual neurons, their coherence is captured only by population-level metrics. One of our contributions in this paper is to promote the use of such metrics.

A body of related work that has received a great deal of attention goes by the name *communication through coherence* (Fries 2005, 2015). Coherence here is similar to what we call synchrony and correlations in the present paper, and is also closely related to rhythmic activity. There have also been a number of theoretical studies focusing on the transference of information through oscillatory dynamics. While not unrelated, these studies differ from ours in that the gamma-band rhythms in this paper are irregular and episodic, following experimental data for the sensory cortices (Henrie and Shapley 2005). Relations between these works and ours will be expanded on in the Discussion.

This paper is organized as follows: In Section 1, we introduce a family of homogeneously connected models to represent local populations and document some spike-firing properties. In Section 2, we build multi-component networks out of the local populations in Section 1 and present the formal definitions of population-level correlations and response times, definitions that will be used throughout. Section 3 studies in some detail correlations between directly connected source and target populations, and Section 4 treats more complicated network motifs. Section 5 is on Methods.

### 1 Single-population models and their spiking patterns

In this section we describe the models for the local populations used in the correlation studies to follow. We also document their gamma-band activity. As we will show later on, these rhythms serve to synchronize connected populations.

#### 1.1 Model description

Local populations in this paper are intended to model local circuits in the cerebral cortex. Typically they consist of a few hundred neurons, three quarters of which are Excitatory (E) and the rest Inhibitory (I). The neurons are randomly and homogeneously connected to one another according to certain specified probabilities, and they are modeled as integrate-and-fire neurons. Similar models were used in Chariker and Young (2015).

The local population models used for the simulations in this paper are networks consisting of 300 E and 100 I-neurons. On average each E-neuron is postsynaptic to about 80 other E-neurons and 50 I-neurons, and each I-neuron is postsynaptic to about 240 E-neurons and 50 other I-neurons. Connectivity between pairs of neurons are subject to variance, and are randomly drawn according to the means above. These connection probabilities, with E-to-E being more sparse than connections that involve I-neurons, are consistent with neuroanatomy.

Leaving details to Supplementary Information, we describe the dynamical interaction within the local population. The dynamics of individual neurons are governed by standard conductance-based leaky integrate-and-fire (LIF) equations, in which the membrane potential *V* of a neuron is normalized so that when it reaches 1, the neuron fires an action potential following which its membrane potential is immediately reset to 0, where it remains in refractory for a couple of milliseconds.

The LIF equations contain 8 parameters. Four of them, *S*^*QQ′*^, *Q, Q*^*′*^ ∈ {*E, I*}, represent the synaptic coupling weights from neurons of type *Q*^*′*^ to neurons of type *Q*. Two others, *τ*_*E*_ and *τ*_*I*_, denote the rates at which the E and I-conductances, which elevate upon the arrival of a spike, decay to zero.

In addition to the synaptic input received from within the local population, each neuron receives an excitatory external drive modeled as a Poisson point process. This drive is independent from neuron to neuron. The external drive has two components: a *synaptic component*, which consists of E-spikes representing input from other regions of the brain with synaptic weight *S*^*QE*^ for neurons of type *Q*, and an “*ambient*” component with a smaller synaptic weight meant to represent all neurotransmitters not specifically modeled. The ambient component’s Poisson rate is assumed to be constant, whereas the synaptic component’s rates, *λ*_*E*_ and *λ*_*I*_ for E and I-neurons, are assumed to be low in background and to increase with drive.

The parameters above are chosen with guidance from realistic models of the visual cortex such as that in Chariker et al. (2016). Their exact values are unimportant for purposes of the present study, as long as they produce reasonable dynamics. For example, since we are primarily interested in correlations between populations when driven by a stimulus, we use values of *λ*_*E*_ and *λ*_*I*_ that produce average E-firing rates of about 15 spikes/sec, which is consistent with the average stimulus-driven firing rates of local populations in realistic situations (Chariker et al. 2016, 2018).

This completes our description of the model; details of the LIF equations and exact parameters used are given in Supplemental Information.

#### 1.2 Gamma-band activity: an emergent phenomenon

Interspike intervals of individual neurons have long tail distributions that have been described as being exponential (Ostojic 2011) or obeying power laws (Baddeley et al. 1997), and Figure 1a shows that our model exhibits a similar behavior. These long tails suggest that individual neurons do not spike rhythmically. To contrast with that, we show in Figure 1b, rasters of the population over a time interval of 500 ms. Here one observes a tendency for the spikes to occur in clusters, leading to rises and falls in firing rates that produce a rhythm in the gamma band, a phenomenon well known to occur in many parts of the brain (Henrie and Shapley 2005; Pesaran et al. 2002; Khawaja et al. 2009; Buzsáki 2011; Chariker et al. 2018). This rhythm cannot be detected by observing individual or even a handful of neurons. It is an example of an *emergent* phenomenon, i.e., a phenomenon that is not part of the rules of operation of individual neurons, but that occurs only as a result of the interaction among neurons.

**Fig 1:**
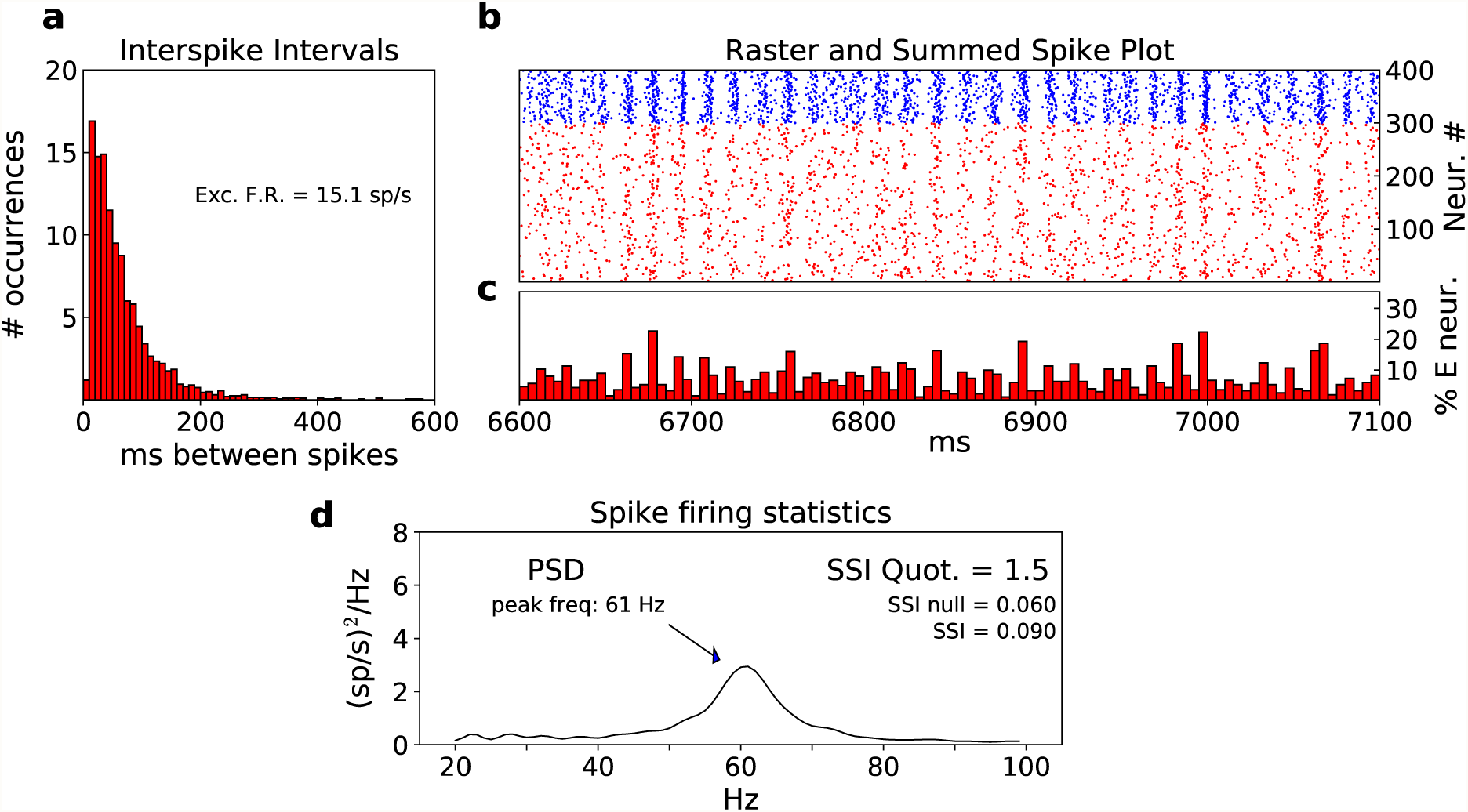
Spiking statistics in a strongly driven regime. **a** Interspike interval plots showing spike firing statistics of individual neurons. We randomly chose 20 E neurons and for each, calculated a histogram of time between each spike with 10 ms bins. We then averaged the 20 histograms to obtain a typical interspike interval plot for an E neuron in the simulation. This was calculated using 8 seconds of simulation data. Simulations using a single neuron produced very similar outputs when sampled over a much longer time period. **b** Population statistics: Raster showing half a second of the simulation. Red dots represent excitatory spikes and blue are inhibitory spikes. A rhythm in the gamma band is clearly visible. **c** Summed spike plot for the E neurons of plot B, showing the fraction of the E-population spiking within each time bin of 5 ms. **d** SSI values and PSD for the regime in panels a, b, and c

To reconcile the seemingly incompatible behaviors shown in Panels a and b of Figure 1, let us first recall how gamma-band rhythms come about. Up until recently, a widely accepted explanation was PING (Whittington et al. 2000), which is the result of a steady external drive together with E-to-I and I-to-E interactions within the local population. PING produces population spikes that are periodic and highly regular, contrary to experimental observations of gamma-band activity in cortex. A more realistic phenomenon called *multiple firing events* (MFEs) was proposed in Rangan and Young (2013), subsequently studied in Chariker and Young (2015), and re-examined in Chariker et al. (2018) using a previously constructed realistic model of the visual cortex. A mechanism for the formation of MFEs is *recurrent-excitation-inhibition* (REI), which works as follows: The crossing of threshold by a few E-neurons leads to recurrent excitation, which may or may not cause other neurons to spike. If it leads to elevated spiking activity, both E and I-neurons will be activated, and the firing event will continue (usually for 2 − 3 milliseconds) until the voltages of the neurons are pushed back by the inhibition. The decay of I-conductance and the depolarization of E-cells due to external input may then lead to the next event. The time constant for I-conductance decay in the LIF equations places the frequency of these spiking events in the gamma band.

A major difference between REI and PING is that in REI, events are not always identifiable, and when they are, they involve variable fractions of the population (depending on the voltages of neurons postsynaptic to the first ones to cross threshold), and inter-event times are variable (depending on the fraction of I-neurons involved). Figure 1c shows that typically no more than 10-15% of the E-population participates in each event. Comparing Panels a-c in Figure 1, one deduces from the firing rates of individual neurons and the gamma frequencies that each neuron participates in only a fraction of the MFEs, skipping over seemingly random stretches of spiking events as the long tail of its ISIs would suggest. This, together with the variations in spiking events, is what makes it possible to have the plots in both Figure 1a and b. The periodic population spikes produced by PING cannot be reconciled with the ISI plots in Figure 1a.

As mentioned in the Introduction, gamma-band rhythms play a role in shaping correlations between brain regions.

##### Quantifying synchrony

Two metrics called *SSI* (spike synchrony index) and *SSI-quotient* are used to quantify the degree of synchrony in a local population. These metrics were introduced in Chariker et al. (2018); we recall them in Section 5 (Methods) for the convenience of the reader.

Informally, the definition of SSI is as follows. We fix a window of length *w* (*w* = 4 ms is suitable for capturing gamma-band activity and is assumed in our simulations). At each E spike fired, we measure the fraction of the E population spiking within a time interval of length *w* centered at the time of this spike, and average over all the E-spikes fired by the population, without distinguishing between the activity of individual neurons. Roughly speaking, SSI= 0.1 means that on average 10% of the excitatory neurons in the population “fire together”, (meaning within the same 4 ms interval); SSI= 1 means the E-population spikes fully synchronously.

SSI-null is the SSI of a perfectly homogeneously spiking population having the same firing rate as the one in question, and SSI-quotient = SSI / SSI-null. For example, SSI-quotient = 2 means that on average, spikes tend to form clusters with twice the density of a uniform distribution.

A more traditional way to quantify both mean frequency and the breadth of the frequency band in gamma activity is to compute its *power spectral density* (PSD). The precise definition of PSD is also recalled in Section 5 for the convenience of the reader.

Figure 1d shows the spike synchrony indices and PSD of the same regime in Panels a-c.

#### 1.3 Regimes with different degrees of synchrony

Gamma-band activity in sensory cortices is typically not much more synchronized than that in the regime shown in Fig 1; our parameters were in fact chosen to emulate data from the visual cortex; see Henrie and Shapley (2005). Different circumstances may produce regimes in other parts of real cortex that are more synchronized, such as at the onset of a stimulus presentation, increased attention, or the effects of drugs (such as anesthesia or ketamine). It is far beyond the scope of the present paper to explore these real cortical phenomena, but it has been suggested (Fries et al. 2001; Fries 2005, 2015) that increased synchrony enhances communication, and we will want to investigate this hypothesis in the context of our spiking neuron models.

We present here systematic ways to produce models with different degrees of synchrony. Similar techniques were used in Chariker et al. (2018). Essentially, we reduce the rise times of postsynaptic excitatory conductances on E-cells, modeled here as a brief time delay, and increase the inhibitory conductance decay times for all neurons. The first has the effect of increasing the speed of recurrent excitation, thereby allowing more E-neurons to participate in a spiking event before it is stopped by I-suppression. The second has the effect of having the suppression last longer, thereby creating a longer period with low firing, and hence, a more synchronized regime.

Panels a and b in Figure 2 show two examples of local populations exhibiting more synchronized spiking dynamics than those in Figure 1, and the regime in Figure 2b is more synchronized than the one in Figure 2a, as can be seen by their SSI-quotients and PSDs. All three regimes, the one in Figure 1 and the two in Figure 2a, b, have similar firing rates.

**Fig 2:**
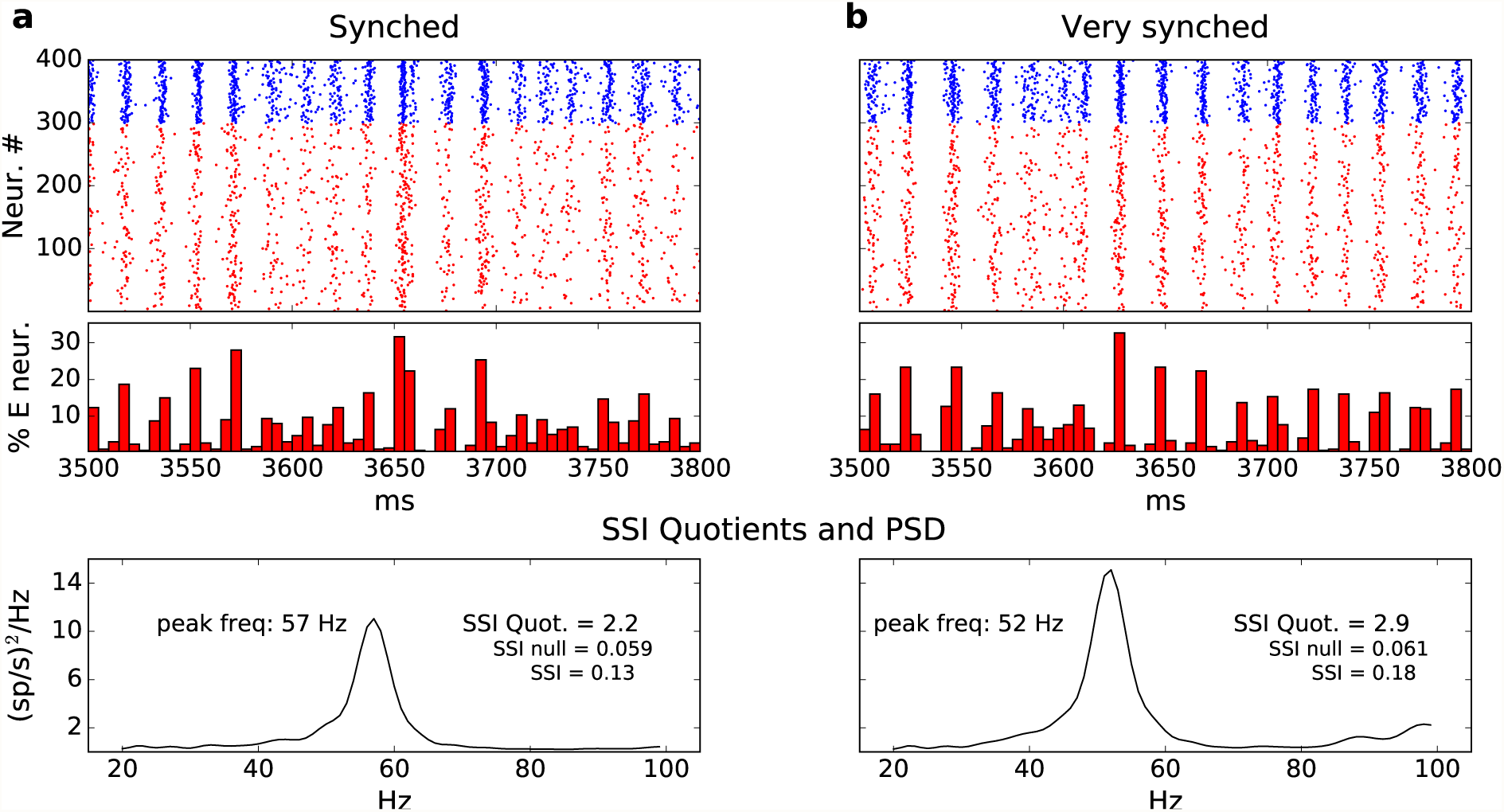
**a** Population statistics for a more synchronized regime. From top to bottom: raster of the simulation, summed spike plot of the E neurons, and SSI and PSD values for the simulation. See Methods for details on computation. **b** Population statistics for a very synchronized regime. This panel follows the format of panel a. Notice the changes in spectral power and peak frequencies as we go from the normal regime in Figure 1d to the synched and the very synched regimes

Networks constructed using parameters from these three regimes are used for the simulations in the rest of this paper. We will refer to the regime in Figure 1 as “normal”, the one in Figure 2a as “synched”, and the one in Figure 2b as “very synched”. We point out that even in the very synched regime, gamma-band activity degrades and resynchronizes (as it does in real cortex) and the PSD is very far from a delta function, which is equivalent to the regime being far from periodic.

### 2 Multi-component Networks and Correlations in Population Activity

The two goals of this section are as follows: One is to introduce the multi-component network models studied in the rest of this paper (Sect. 2.1), and the other is to fix a notion of correlations in spiking activity between populations (Sect. 2.2).

#### 2.1 Multi-component network models

##### The simple feedforward case

Given two networks, 𝒩_1_ and 𝒩_2_, of the type in Sect. 1.1, and a number *p* ∈ (0, 1), we first describe how to construct a simple feedforward network, henceforth abbreviated as

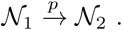

Here, 𝒩_1_ is the source network, 𝒩_2_ is the target network, and there are excitatory connections from 𝒩_1_ to both E and I-neurons in 𝒩_2_, chosen such that on average, each neuron in 𝒩_2_ receives a fraction *p* of its total excitatory input from 𝒩_1_. “Total excitatory input” includes all sources that increase E-conductances: from E-neurons within 𝒩_2_, from the two external sources described in Sect. 1.1, and from neurons in 𝒩_1_. Presynaptic neurons from 𝒩_1_ are chosen randomly.

To build this composite network, we simply take away a suitable fraction of the synaptic component of the external drive for neurons in 𝒩_2_, and replace it by spikes from 𝒩_1_. Consider for definiteness, the E-neurons in 𝒩_2_. Using the mean E-firing rate of 𝒩_2_, the mean number of presynaptic E-cells, synaptic coupling weights, and external drive rates, we compute the number of spikes, *x*, that provide a fraction *p* of their total excitatory input. We then decrease the synaptic component of the external drive by *x* spikes/sec, and estimate, based on the E-firing rate in 𝒩_1_, the number *k* of presynaptic E-neurons from 𝒩_1_ that will produce *x* spikes. For each E-cell in 𝒩_2_, we then randomly draw a group of E-cells in 𝒩_1_ of size *k* in mean, and assign connections from this group to the cell in question.

The same algorithm applies to I-neurons in 𝒩_2_. The fraction of current, *p*, will be referred to in the future as the “connectivity” from 𝒩_1_ to 𝒩_2_. For the spikes from 𝒩_1_ to 𝒩_2_, we assume an additional *δ* ms transmission time. Unless stated otherwise, *δ* = 1 in our simulations. Detailed formulas for the construction of 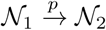 are given in Supplementary Information.

##### Building general multi-component networks

The coarse architecture of a multi-component network consists of a finite, directed graph, i.e., a set of nodes denoted {1, 2, …, *n*_0_}, a collection of directed edges {*i* → *j*} with 1 ≤ *i, j* ≤ *n*_0_ and *i* ≠ *j*, and a collection of numbers {*p*_*ij*_}, one for each edge. We can now construct a multiple-component network where each node *i* in the graph corresponds to a single-population model of the kind in Section 1, each directed edge {*i* → *j*} represents the presence of a connection from component *i* to component *j* with connectivity *p*_*ij*_. We will use this construction for the larger network motifs used in section 4.

#### 2.2 Correlations between populations and response times

In this paper, we focus on correlations on brief time-scales of a few, up to ∼ 10, milliseconds (ms). These timescales are of interest for brain regions separated by no more than two or three synapses.

Given two networks 𝒩_1_ and 𝒩_2_ that are directly or indirectly connected (or not connected at all), we now give the formal definition of correlations between populations as used in the rest of the paper. As we have stressed throughout, this is strictly a population-level metric. It can be extended to correlations between smaller samples of neurons randomly drawn from 𝒩_1_ and 𝒩_2_, but we leave that for later.

The (unadjusted) notion of correlation of interest to us is the time correlation of the spiking activity in the two local populations 𝒩_1_ and 𝒩_2_. We fix a large time interval [0, *T*], and let *F*_1_(*t*) and *F*_2_(*t*), *t* ∈ [0, *T*], be the instantaneous population firing rates of 𝒩_1_ and 𝒩_2_ (to be made precise momentarily). Then the quantity of interest is

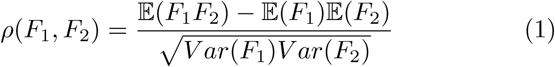

where the underlying probability is the uniform distribution on [0, *T*].

The precise meaning of “instantaneous population firing rate” of 𝒩_*i*_, *i* = 1, 2, is as follows: For *t* = 0, 1, 2, …, *T* − 4 ms, we let

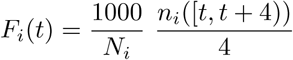

where *n*_*i*_([*t, t* + 4)) is the total number of spikes fired by all neurons in 𝒩_*i*_ on the time interval [*t, t* + 4) and *𝒩*_*i*_ is the total number of E-neurons in 𝒩_*i*_ (300, as per Section 1.1). The choice of 4 ms above reflects the time scales we find relevant for our purposes: we are not especially interested in pinpointing the exact timing of spikes, but want short enough time intervals to reflect gamma-band activity, which dominate local-in-time firing patterns. To smooth out the statistics further, we use sliding windows.

We will refer to the definition of correlation above as “correlation without adjustment”. Under many conditions, correlations in spiking activity between two populations are larger if one measures the spike times of one network with a time delay, i.e., if instead of *ρ*(*F*_1_, *F*_2_), we consider

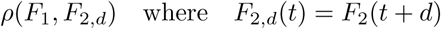

for a suitable *d*.

Consider, for example, the network 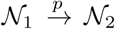 discussed at the beginning of Sect. 2.1. It is reasonable to expect that the spiking events in 𝒩_1_ would match those in 𝒩_2_ better if we measure the spike times in 𝒩_2_ with a delay. To locate the optimal time delay, i.e., the time delay that maximizes their correlations (if there is one), we computed *ρ*(*F*_1_, *F*_2,*d*_) for various values of *d*, at 0.5 increments for *d* from 0 to 30 ms. Figure 3a shows the function *d* 1↦ *ρ*(*F*_1_, *F*_2,*d*_) for *p* = .075, i.e., when 7.5% of the excitatory drive in 𝒩_2_ comes from 𝒩_1_. (Values of *p* in the real cortex obviously vary, but *p* ∼ 0.075 to 0.1 is thought to be fairly typical.) The 5 graphs superimposed are for 5 different networks (constructed with the same connection probabilities) and 5 different sets of initial conditions. Across the trials, the optimal time delay is approximately 3.5 − 4 ms. The low trial-to-trial variation of these plots suggests that this optimal time delay is intrinsic to the network architecture and is not seriously affected by network details or initial conditions that vary trial to trial.

**Fig 3:**
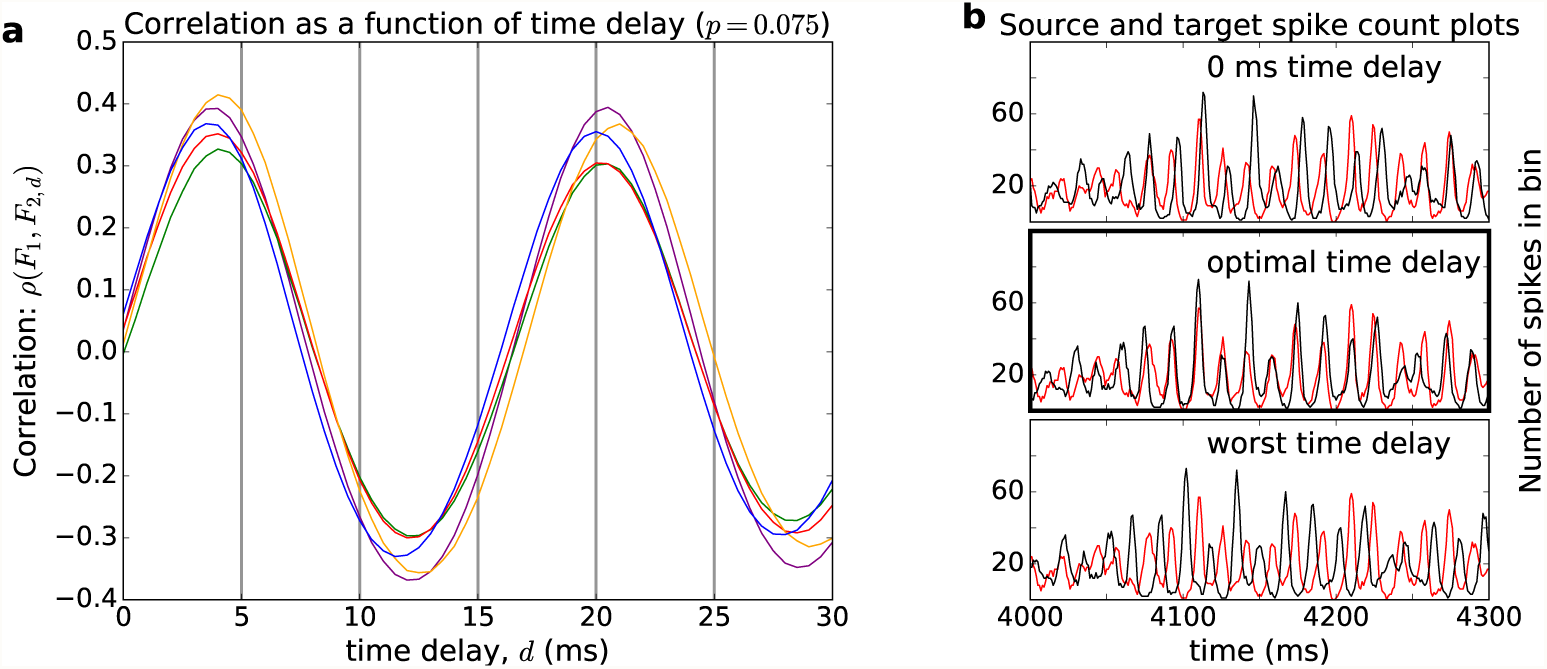
Correlations with time delay in network activity for 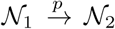, *p* = 0.075. **a** Computed are correlations between *F*_1_(*t*) and *F*_2_(*t* + *d*) as functions of *d* (for 0.5 ms increments of *d*) for two networks connected as in Sect. 2.1. Each color represents a different trial using a different network with the same connection probabilities. The locations of the first peaks, which occur at approximately 3.5 − 4 ms for all the trials, are taken to be the response time of 𝒩_2_ to 𝒩_1_. **b** shows the superimposed plots of *F*_1_(*t*) and *F*_2_(*t* + *d*) for one of the trials in panel a, for *d* = 0, 3.5 and 11.5 ms from top to bottom, respectively; the graph of *F*_1_(*t*) is in red, and that of *F*_2_(*t* + *d*) is in black. Observe that the two plots in the bottom graph of panel b are near anti-phase, consistent with the location of the first minimum in panel a

Observe that the 3.5 − 4 ms optimal time delay is significantly longer than the 1 ms transmission time imposed. This is because spikes from 𝒩_1_ do not cause spikes in 𝒩_2_ immediately. Instead, they raise the excitatory conductance for neurons in 𝒩_2_ and bring their membrane potentials closer to threshold. During this process, some E-neurons in 𝒩_2_ may fire. This will further raise the membrane potential of other neurons in the network, possibly pushing a larger percentage of the E- and I-neurons in 𝒩_2_ to spike, and subsequently causing an MFE; see the discussion in Sect. 1.2. As this is not an instantaneous process, allowing a time delay for the activity in 𝒩_2_ to build up increased the correlations between *F*_1_ and *F*_2_.

Figure 3b shows the functions *F*_1_(*t*) and *F*_2_(*t* +*d*) as functions of *t* for *d* = 0, 3.5, 11.5, from top to bottom, respectively. In the top plot, one sees that for *d* = 0, i.e., without a time delay, network activity in 𝒩_1_ (red) peaks a little ahead of that in 𝒩_2_ most of the time. In the middle plot where the delay is optimal, one sees, in the center half of the plot, an excellent alignment of the MFEs produced by 𝒩_1_ and 𝒩_2_. Such alignments are not always present, however, because gamma rhythms degrade from time to time, as can be seen at the beginning of the time interval shown. In the bottom plot, one notes that the rhythms of the two networks are almost exactly anti-phase – not unexpected given that correlation is at its most negative with a delay *d* = 11.5 ms; see Figure 3a.

In the rest of this paper, when there exists a number *d* such that *independent of initial conditions*, the correlations between two networks, 𝒩_1_ and 𝒩_2_, are maximized when the spike times of 𝒩_2_ are measured with a delay of ≈ *d* ms, we will call *d* the *response time* of 𝒩_2_ to 𝒩_1_, and refer to the correlations with this delay as the *peak correlations* between the two networks. We remark that when a response time exists, it represents an intrinsic relationship between the two networks, namely how long it takes for spiking events in one to “cause” spiking events in the other. In the case of 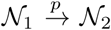 above, response time is well-defined. For two networks embedded in a more complex multi-component network, it is less clear if there is a well-defined notion of response time.

The presence of a response time or phase-shift between post-synaptic and pre-synaptic firing has been observed experimentally (Zandvakili and Kohn 2015; Fries 2015, 2005; Jia et al. 2013; Bastos et al. 2015). On the other hand, we are unaware of in-depth theoretical studies of this issue.

### 3 Analysis of two-component networks with feedforward drives

This section studies exclusively the two-component network

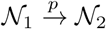

where 𝒩_1_, 𝒩_2_, and *p* are as defined in Sect. 1.1. In Sect. 3.1 we investigate correlation properties as functions of *p* and degree of synchrony, and in Sect. 3.2, we study the dependence of correlations on sample size, connecting population-level metrics to correlations between pairs of neurons.

#### 3.1 Correlations as functions of connectivity and synchrony

The results of this subsection are summarized in Figure 4. We identify below a few points of note, offering mechanistic explanations when we can.

**Fig 4:**
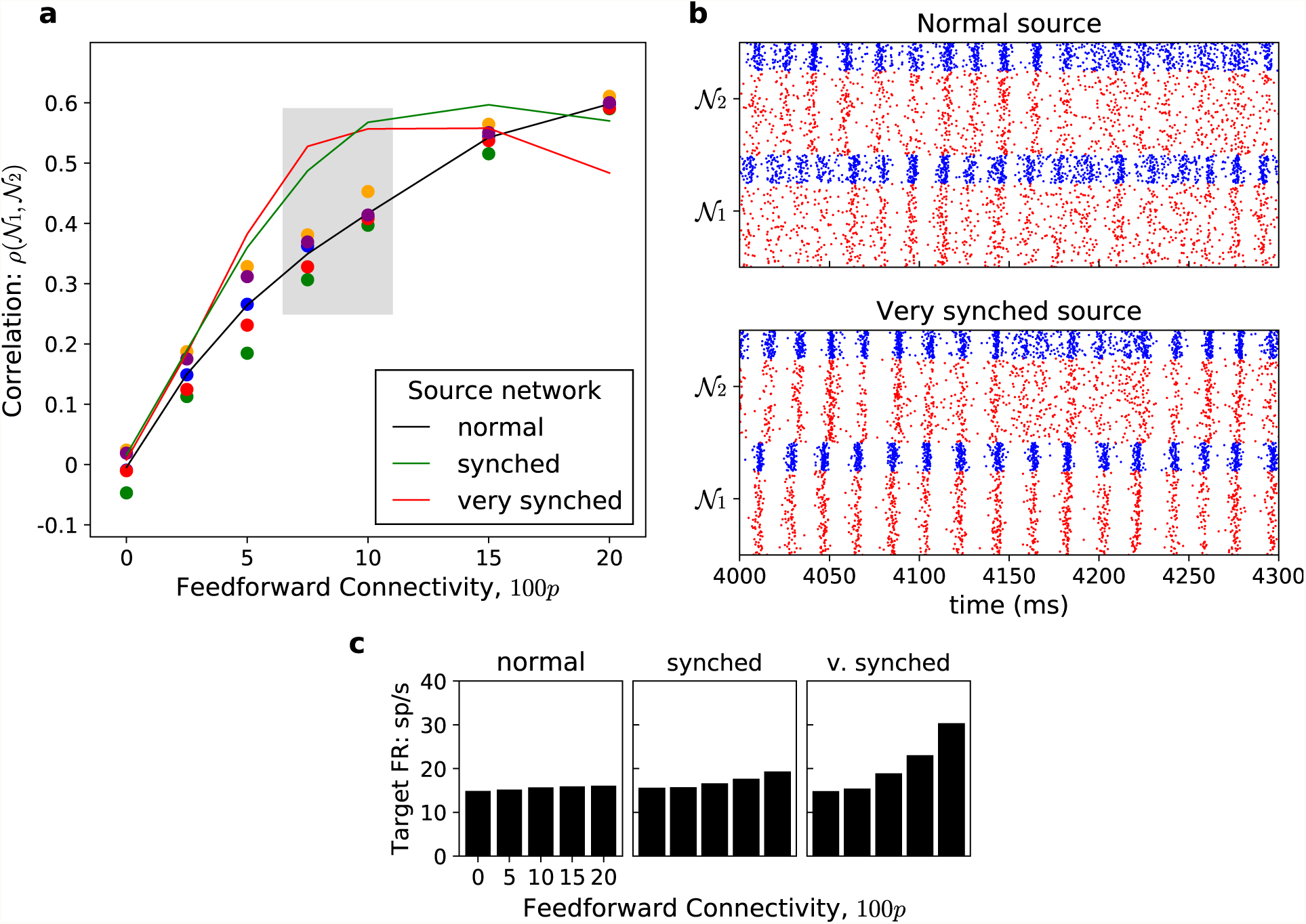
Correlations between source and target networks. All target networks are in the normal regime as defined at the end of Sect. 1. **a** Correlation as function of connectivity. Shown are peak correlations as function of *p*. The three different plots represent different source regimes. For the black line representing a normal source, the color dots are results from 5 trials using 5 different networks drawn with the same parameters. Note the strong correlations, as well as the low trial-to-trial variability. The values for the other lines were also averaged over 5 trials. Note that for the range that is most realistic (grey box), the more synchronized source regimes produce higher correlations. **b** Rasters of 𝒩_1_ and 𝒩_2_ with a normal source (top panel) and a very synched source regime (bottom panel). **c** Mean firing rates of 𝒩_2_, the target network. *x*-axis is the percentage connectivity, 100*p*. The source networks are, from left to right: normal, synched, and very synched. Bar graphs show the average of 5 trials, 8 sec each. Note that for *p* ≤ 0.1, firing rate did not increase appreciably with the increased synchrony of the source network

First we discuss the black curve in Panel a of Figure 4, showing peak correlations between two networks in normal regimes connected at various values of *p*. For each value of *p*, five different trials using independently drawn networks (with the same connection probabilities) and different initial conditions were performed. The variation from trial to trial was small, so the metric is quite robust. Response times were well defined and easy to locate down to *p* = 0.05. While it is to be expected that correlations will increase with *p*, it is surprising how large these values are: For *p* = 0.075 to 0.1, i.e., with no more than 10% of the excitatory input to neurons in 𝒩_2_ coming from 𝒩_1_, peak correlations can be as large as 0.4. Below, we provide an explanation for this unexpected finding.

##### Analysis

As we have seen in Figure 3b (middle panel), the strong positive correlation between 𝒩_1_ and 𝒩_2_ comes from the tendency for their gamma rhythms to align. In other words, *gamma-band activity helps synchronize the two networks.* This may at first seem contrary to our discussion in Sect. 1.2, where we described how gamma rhythms are generated within local populations: How can two rhythms that are locally, hence independently, generated match up so well irrespective of initial condition? A more careful examination revealed that there is no contradiction at all: while the tendency to produce a rhythm comes from the internal dynamics within a local population, there is *no intrinsic timing* to the peaks and valleys of the spiking events. A little extra help from 𝒩_1_ facilitates events in 𝒩_2_, increasing the likelihood that the rhythms will line up. However, since these rhythms degrade, they also become “unsynchronized” after a few events, hence the correlations are not higher than they are.

This raises a number of interesting questions: How do correlations depend on the degree of synchrony of the networks, and must 𝒩_2_ share the peak gamma frequency of 𝒩_1_ for its gamma rhythm to lock onto the one produced by 𝒩_1_? The other plots in Fig 4a, answered the first question: *Synchronous source networks produce higher correlations, i.e., they are more effective in entraining populations downstream*. This is true with or without the peak gamma frequencies of the source and target regimes coinciding.

For example, when we used a very synched network, with a peak gamma frequency of ∼ 52 Hz, to drive one in the normal regime, with peak gamma frequency > 60 Hz, correlations went up to 0.5 − 0.6. This is considerably higher than the correlations for normal-driving-normal networks at corresponding values of *p*. It may well be true, however, that all other things being equal, correlations are higher when the intrinsic frequencies of the gamma rhythms in the two networks coincide, as in the case of a very synched source driving a very synched target.

As to why synched source networks are more effective in eliciting a higher correlation with the target network, our analysis suggests the following mechanistic explanation:

##### Analysis

In a synched 𝒩_1_, the spiking events are more clearly defined, with cleaner gaps in between. The more convergent excitatory input provides a stronger drive to 𝒩_2_ during MFEs, and a significantly decreased drive between MFEs. This causes 𝒩_2_, which is affected by this time-varying nature of the feedforward input, to lock onto the peaks and troughs of 𝒩_1_-activity (with a phase-shift). Hence, we see an increase in correlation compared to where there is a more ambiguous feed-forward signal from an 𝒩_1_ in the normal regime. Incompatible gamma frequencies may lower correlation but is not a serious obstruction because gamma rhythms degrade and resynchronize, so 𝒩_2_ can realign itself with 𝒩_1_ again after several spiking events.

These findings are consistent with the communication through coherence ideas in Fries (2005), provided better communication is interpreted to mean higher correlations. There is another commonly held belief, possibly (but not necessarily) suggested also in Fries (2005), that synchronized networks are more effective in causing higher firing rates downstream. We investigated the issue, and found that for *p* ≤ 0.1, synchrony in the driving network does not lead to increased firing rate in the target network. Such increases are observed only for driving networks that are very synched, and for *p* large, neither of which is typical. These results are illustrated in Figure 4c.

#### 3.2 Correlations between smaller samples

Correlations between pairs of neurons have been studied in detail experimentally, and the numbers found have been very small (Zandvakili and Kohn 2015; Kuhn et al. 2003). Since electrophysiology does not offer information on the simultaneous activity of large numbers of neurons, and calcium dynamics are too slow to capture correlations on these short time scales, population level correlations of the kind we have been studying are, at least for now, inaccessible in the laboratory. Correlations between smaller samples, however, can be captured by multi-unit recording. Hence, a natural progression would be to understand theoretically how correlations depend on sample size, such as the minimum number of E-neurons needed to get a reasonable estimate of the population correlation and optimal time delay.

We studied this in the 𝒩_1_ → 𝒩_2_ network using samples of *𝒩* E-neurons from each layer, where *𝒩* varied from 1 to the size of the full population. The correlation between a set of *𝒩* E-neurons from 𝒩_1_ and a set of *𝒩* E-neurons from 𝒩_2_ is as defined in Sect. 2.2, but with *F*_1_ and *F*_2_ defined using only the spikes times of the 2*N* neurons in the samples;.

Results for *p* = 0.075 are shown in Figure 5, with 5 trials for each *N*. Here *N*= 1 corresponds to correlations between pairs of neurons. We see that the numbers are very small, about a tenth of the population correlations. In addition to having small correlations, the trial-to-trial variances are large, having the same order of magnitude as the correlations themselves. These observations are consistent with individual neuron spikes being almost “random”, as depicted in Figure 1a. We also see that as *N* increases from a small value, correlations increase rapidly, stabilizing at about *N*= 100 and asymptoting eventually to the population value.

**Fig 5:**
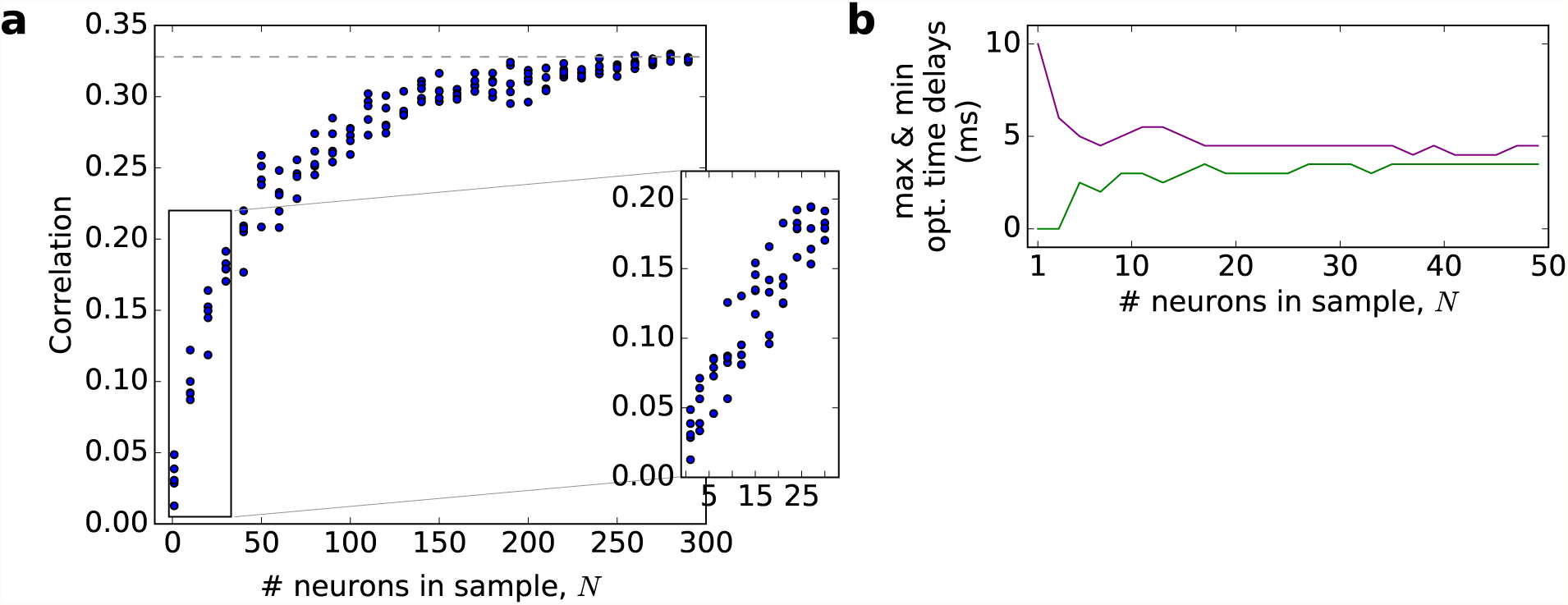
Correlations and optimal delays as functions of sample size *N* **a** Correlation between target and source (both normal regime, *p* = .075) as *N*increases. Results are based off of one, 8 second long simulation, with correlations computed for *N* = 1, 10, 20, 30, …, 290 randomly selected E-neurons. Shown are results of 5 trials (a different sample for each trial) for each *N*. The population correlation using all 300 neurons is about 0.33 and is shown by the gray, dashed line. The inset shows more detail for smaller samples, with *N* = 1, 3, 6, 9, …, 30. **b** Maximum and minimum optimal time delays as a function of *N*, for the same simulation in panel a. The range of time delays was computed over 10 trials (again using a different sample of neurons in each trial)

A noteworthy observation is that when using as few as 10 neurons from each layer, the optimal time delays already stabilize and are around 3.5-4 ms, which is the response time discussed above in Section 2.2. Panel b in Figure 5 shows the range of optimal time delays narrowing rapidly and stabilizing fairly quickly as *N* increases. Thus unlike population-level correlations, which are strongly dependent on sample size, response time measurements require few neurons and are relatively accessible using present technology.

### 4 Correlations in Larger Networks: Five Canonical Motifs

Brain regions are, in general, interconnected in complicated ways; see e.g. Binzegger et al. (2009). Consider, for example, the primary visual cortex V1, which has 6 layers with further subdivisions within layers. Diagrams showing the connectivities among layers can be complicated (Sincich and Horton 2005). Input from different regions converge, there are feedback loops, and there can be multiple pathways for signals to travel from Point A to Point B. In this section, we study the correlations and response times between pairs of components in a few canonical network motifs that are known to occur in the real cortex. Knowledge of such characteristics in these smaller multi-component networks can help dissect how spiking in one area of the brain affects spiking in another, when these components are embedded in much more complex distributed networks.

As has been noted in Sects. 2.2 and 3.1, positive correlations between populations arise in part from the alignment of their gamma rhythms. We reiterate that gamma rhythms in our local populations (Sect. 1.2) are irregular; they degrade and resynchronize as in real cortex and are quite far from truly periodic and/or completely synchronous. These characteristics are relevant to some of the phenomena below.

All correlations below refer to those between entire local populations. Each network motif was selected to answer a question or test a hypothesis, and the result will contain a specific message.

#### Motif A. Feedforward chain

We consider here a chain of the form

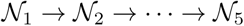

where each 𝒩_*k*_ is a local population described in Section 1.1, following a normal regime. Correlations between 𝒩_1_ and 𝒩_2_ were studied in section 3. We are now interested *ρ*(𝒩_1_,𝒩_*k*_), the correlations between 𝒩_1_ and 𝒩_*k*_ for *k* = 3, 4, 5. Similar feedforward networks have previously been studied (Diesmann et al. 1999; Zandvakili and Kohn 2015).

The two messages from this study are as follows: (1) While correlations between consecutive layers are strong, they decay rapidly with the number of intermediate layers, so that firing patterns further downstream are only weakly correlated to those of the source layer; see Fig 7a. (2) Meaningful response times become fuzzier but persist for 2-3 layers downstream: One can see in Fig 7b that optimal time delays between 𝒩_3_ and 𝒩_1_ are fairly well described by 2*d*; between 𝒩_4_ and 𝒩_1_, 3*d* is a reasonable indicator of optimal time delay in 3 or 4 out of the 5 trials, while the situation between 𝒩_5_ and 𝒩_1_ becomes even less clear.

**Fig 6:**
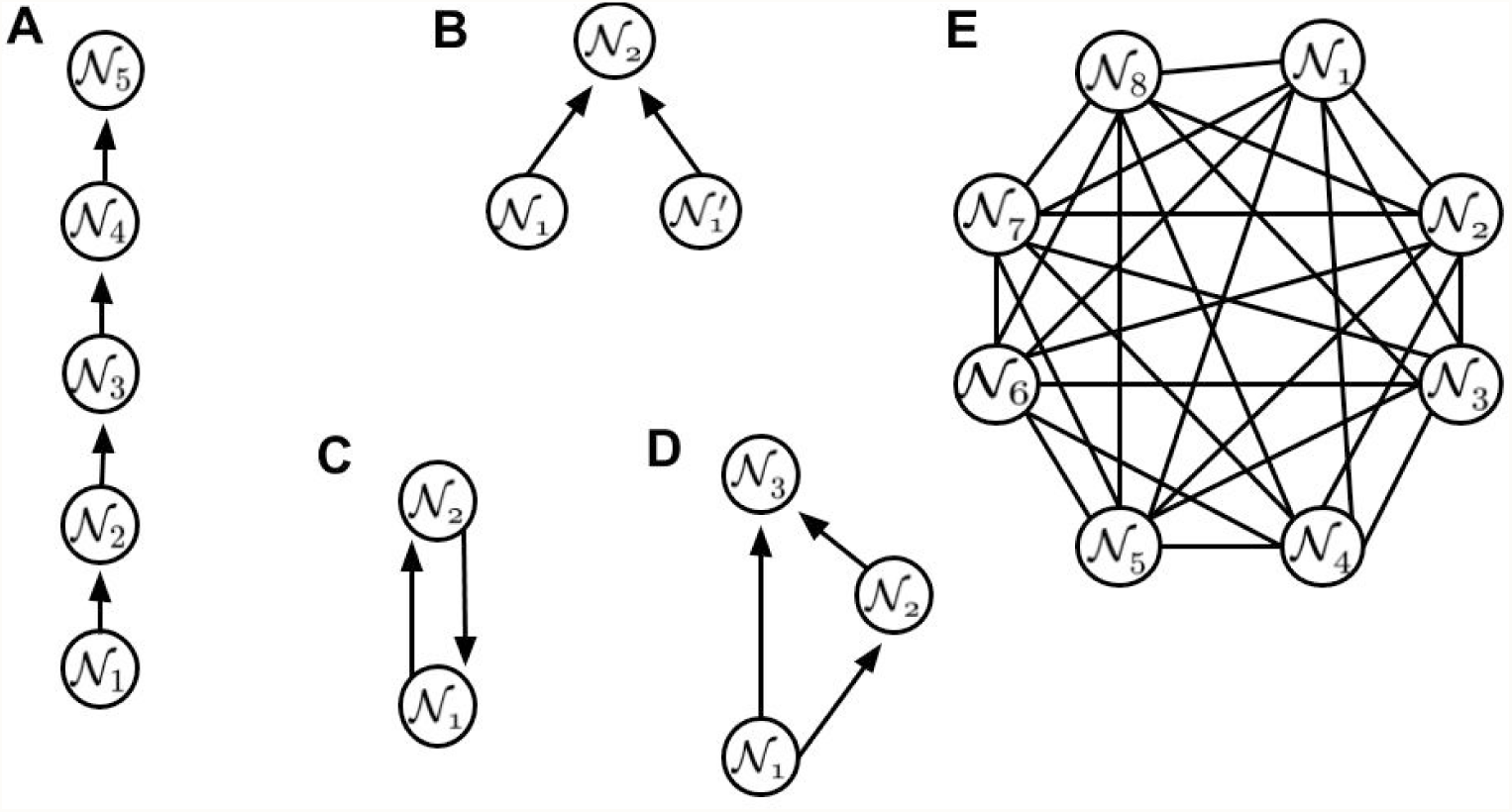
Diagrams of five network motifs, referred to in the text as Motifs A-E

**Fig 7:**
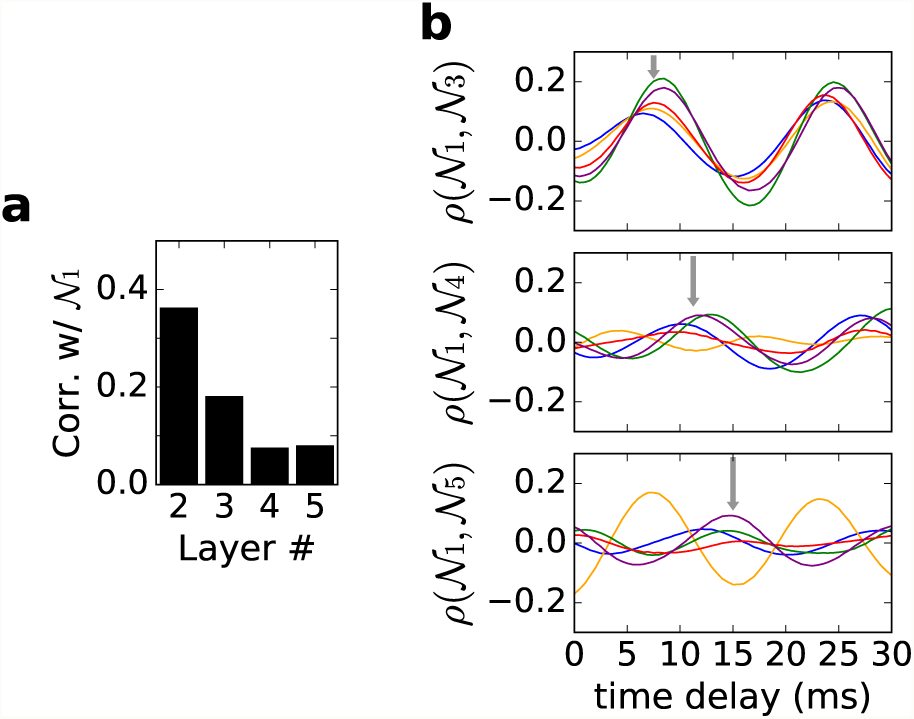
Feedforward chain. All populations follow the normal regime, with 7.5% connectivity from layer to layer. **a** Correlations between 𝒩_1_ and 𝒩_*k*_ for *k* = 2, 3, 4, 5, each averaged over 5 trials. **b** Correlations as a function of time delay for the same 5 trials in panel a. Grey arrows are placed at 2*d,* 3*d* and 4*d* (top to bottom) where *d* is the response time of 𝒩_2_ to 𝒩_1_ from Sect. 2.2. They are to be compared to potential values of response times between 𝒩_1_ and 𝒩_*k*_ for *k* = 3, 4, 5

#### Motif B. Two independent sources

Here we consider a network 𝒩_2_ driven by two local populations

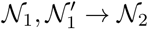

where 𝒩_1_ and 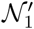 are assumed to be independent: they are unconnected and driven by independent Poisson processes, so there is no reason why the timing of their spiking events would coincide. In this study, we sought to answer the following questions: Comparing this motif to the simple feedfor-ward chain 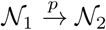, should we expect that “interference” from 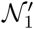 will disrupt the transference of gamma patterns from 𝒩_1_ to 𝒩_2_, thereby lowering their correlation? Will 𝒩_2_ lock on to one of its sources, or try to follow both, and what causes it to prefer one over the other?

We performed simulations with *p* = 0.075, and found that when all 3 networks are in the normal regime, *ρ*(𝒩_1_,𝒩_2_) and 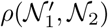 hover around 0.3, which is a little, but not much lower than the correlation of 0.35 for the simple 𝒩_1_ → 𝒩_2_ chain. One of *ρ*(𝒩_1_,𝒩_2_) or 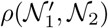 is sometimes a little larger than the other depending on network details; see the scatter-plot in Fig 8a. In the same scatter-plot, we show that when 𝒩_1_, 𝒩_2_ are normal but 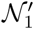 is synched, 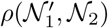 is significantly higher than *ρ*(𝒩_1_,𝒩_2_). This is consistent with our earlier observation (Sect. 3.1) that synchronized networks are more effective in entraining spiking events downstream.

**Fig 8:**
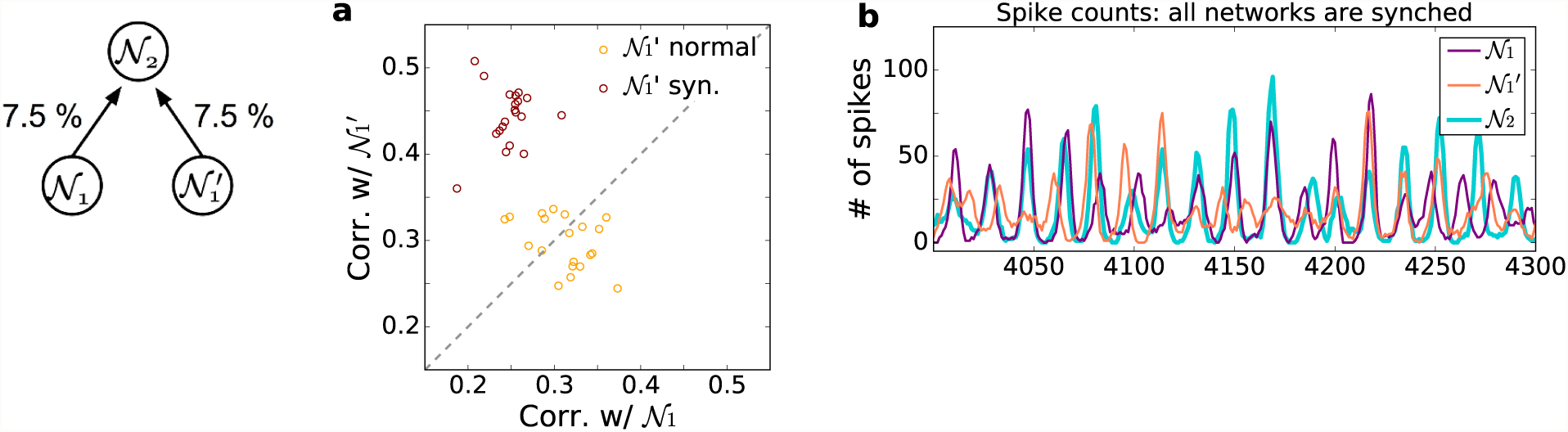
Two independent sources driving one target. **a** Scatterplots showing correlations of 𝒩_2_ with 𝒩_1_ and 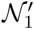, when both 𝒩_1_ and 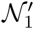 are normal (orange) and when 𝒩_1_ is normal and 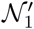 is synched (purple). Results from 20 trials are shown. **b** Illustration of how 𝒩_2_ switches between aligning with 𝒩_1_ and 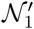. Local-in-time spike rates for 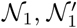, and 𝒩_2_, all synched, are shown on a time interval of 300 ms. Observe that 𝒩_2_ tracked 𝒩_1_ for 30 ms or so before 4050, then slowly shifted to 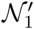 which it tracked for a while, to switch back to 𝒩_1_ by 4130. Sometimes the rhythms of 𝒩_1_ and 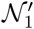 coincide. The plot for 𝒩_2_ has been corrected for response times of ∼ 4 ms

These results raised the following questions: How can 𝒩_2_ align with two separate populations with independent spike times, and why does interference from 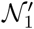 not lower *ρ*(𝒩_1_,𝒩_2_) more?

#### Analysis

We examined more carefully the temporal dynamics of the three networks, and found that 𝒩_2_ follows 𝒩_1_ for a short time period, then aligns itself with 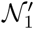, switching back and forth, as illustrated in Figure 8b. In other words, 𝒩_2_ manages to lock on to two networks with independent dynamics by aligning with each one for only a fraction of the time. As to why *ρ*(𝒩_1_,𝒩_2_) was not significantly lowered by the “distraction” from 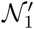, observe that even in the absence of 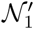, the rhythms of 𝒩_1_ and 𝒩_2_ are only partially aligned (Fig 3b, middle panel): they match up and misalign as each population’s gamma rhythms wax and wane, leaving plenty of room for 𝒩_2_ to engage in other firing patterns.

#### Motif C. Network with feedback

We consider next two-component networks of the form

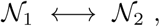

i.e., neurons in 𝒩_2_ both receive excitation from 𝒩_1_ and supply excitation to 𝒩_1_. The question of interest is: will recurrent excitation help synchronize the two populations, raising *ρ*(𝒩_1_,𝒩_2_)? There is a vast literature on the synchronization or phase-locking of coupled oscillators (see e.g. Pikovsky (2001)), and we would like to investigate if the situation here is similar.

Our simulation results were unexpected: We found that with *p* = 0.075 in both coupling directions, the presence of feedback *lowered* correlations significantly from 0.35 to about 0.2, a result we confirmed by varying the feedback connectivity from 0 to 0.075; see Figure 9.

**Fig 9:**
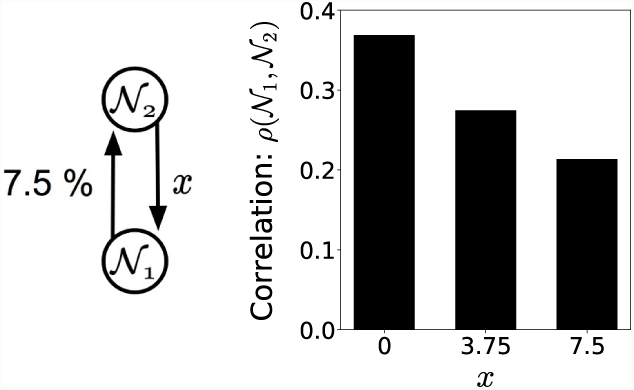
Two-component network with feedback. Both networks are in the normal regime. Feedforward connectivity is fixed at 7.5%, and the bar graph shows correlations, averaged over 5 trials, as function of feedback connectivity

#### Analysis

To explain these findings, we hypothesized that the pulses that originated from 𝒩_1_, passed to 𝒩_2_, and returned to 𝒩_1_ would arrive with a 7-8 ms delay (it takes 3.5-4 ms for 𝒩_2_ to respond and another 3.5-4 ms for 𝒩_1_ to respond to 𝒩_2_). As this was about halfway into 𝒩_1_’s gamma cycle, these pulses were unlikely to precipitate MFEs, i.e., the recurrent excitation was ineffective. Also, this timing was not conducive to the alignment of gamma events in the two networks, which as we have shown is important for positive correlations (see the Analysis in Sect. 3.1).

To test the validity of these conjectures, we performed simulations in which we gradually reduced the rise times of E-conductances and transmission times between 𝒩_1_ and 𝒩_2_ (see Supplementary Information and Sect. 2.1) to speed up the recurrent excitation. We found that as rise and transmission times were decreased to 50% of their original values,

i. in the feedforward chain 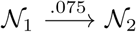 correlations increased from ∼ 0.36 to ∼ 0.46;
ii. with a *p* = .075 feedback, the corresponding values rose steadily from ∼ 0.2 to ∼ 0.6.

These results support the conjecture that recurrent excitation can increase correlations, but the arrival times of spikes, and likely other factors, also play a role.

Unlike the case of a feedforward chain when the target network is influenced by the source network in a relatively straightforward way, feedback creates a more complex dynamic. A prudent conclusion is that the situation warrants a more in-depth study, which we leave to future work.

#### Motif D. Multiple pathways

In the real brain, signals are often passed from Point A to Point B via multiple pathways, some more direct than others. The simplest example of a circuit in which there are two ways to go from one local population to another is shown in Figure 10.

**Fig 10:**
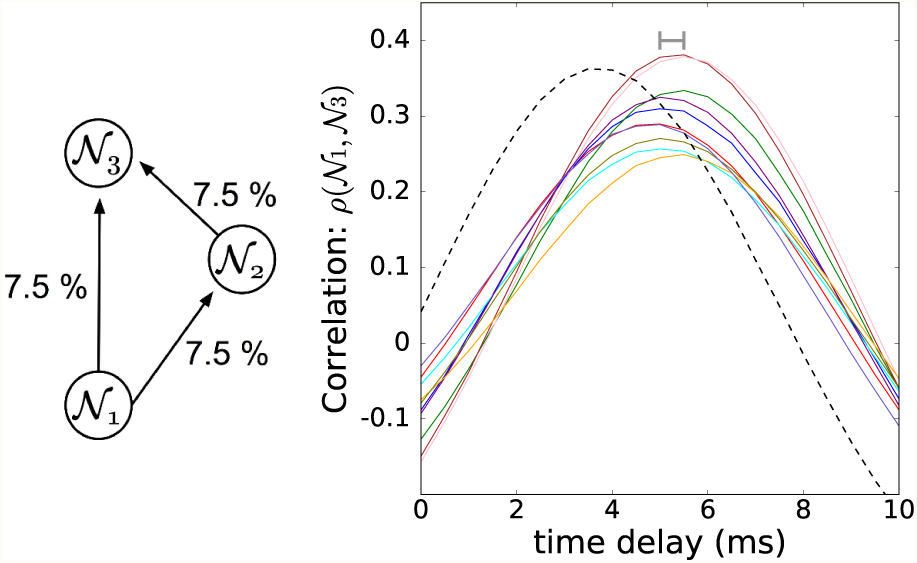
Direct and indirect path from source to target, with *p* = 0.075 for each feedforward connection. The line plots show the correlations between 𝒩_1_ and 𝒩_3_ as functions of time delays for 10 different trials. The network was redrawn and initial conditions changed for each trial. The black, dashed line shows the plot for the simple feedforward chain 𝒩_1_ → 𝒩_2_ case, as in Figure 3a. The horizontal grey bar indicates the spread of optimal time delays across the 10 trials

What intrigued us here was the following: The presence of two different pathways is akin to twice the connectivity from 𝒩_1_ to 𝒩_3_. Since correlations in simple feedforward chains increase rapidly with connectivity (Figure 4a), would the presence of a second pathway lead to higher correlations between 𝒩_1_ and 𝒩_3_, and which pathway would dictate the response time, if it is defined?

Our findings are summarized in Figure 10. Interestingly, there is a fairly well defined response time, which at 5−5.5 ms, is a compromise between the two response times expected for the different paths (see the results for Motif A in Fig. 7b). As for correlations, *ρ*(𝒩_1_,𝒩_3_) is a little lower than that of the simple feedforward chain 𝒩_1_ → 𝒩_2_, but not by too much, as one can see by comparing the solid and dashed graphs in Figure 10.

We hypothesize that as in the feedback case, the differing response times provide interference in internal gamma rhythms, which serves to counter the increased excitation in the presence of multiple pathways going from Point A to Point B.

#### Motif E. A ring network with connectivity decaying with distance

Unlike the previous examples, this motif is intended to mimic networks within a layer of cortex, e.g. layer 2/3 of V1, where in addition to local circuits, long-range excitatory connections exist between domains preferring like orientation in different hypercolumns. These connections can extend 2 − 3 mm, and are expected to become less dense as distance increases. Here we will use a 1D ring of local populations rather than a 2D network (real cortex is more like a 2D surface) because in 2D, the number of local populations involved become large very quickly with radius, making the simulations unwieldy.

The models studied here are densely connected locally, with connectivity decaying with distance as in V1. Transmission times are also assumed to be slower with larger distance, as in V1.

Specifically, we considered a motif with 8 local populations, labeled 𝒩_1_, 𝒩_2_,… 𝒩_8_, to be thought of as arranged in a circle. The distance between 𝒩_*i*_ and 𝒩_*j*_, dist(𝒩_*i*_, 𝒩_*j*_), is defined to be 1 if 𝒩_*i*_ and 𝒩_*j*_ are neighbors, and to be 1+ the number of populations separating 𝒩_*i*_ and 𝒩_*j*_ along the shorter arc of the circle, in general. The connectivities *p*_*ij*_ are taken to be 0.04, 0.02, 0.01, 0.005 for dist(𝒩_*i*_, 𝒩_*j*_) = 1, 2, 3, 4 respectively. Thus local populations in this motif are all-to-all coupled, with coupling strength between populations decreasing exponentially as the distance between them increases. Between 𝒩_*i*_ and 𝒩_*j*_ we impose a synaptic transmission delay of 3 ∗dist(𝒩_*i*_, 𝒩_*j*_) ms. These delays are consistent with the slow transmission times within individual layers of V1, measured to be 5 − 10 ms per mm (Grinvald et al. 1994).

The results are summarized in the scatter-plot in Fig 11. Correlations *ρ*(𝒩_*i*_, 𝒩_*j*_) between all pairs were computed using the optimal time delay for that pair. The overall trends, indicated by the crosses, are that as distance increases, correlations decrease and optimal time delays increase. They are qualitatively similar to those for the feedforward chain in Sect. 4.1. Correlations between neighbors are lower than in most of the previous examples, consistent with the weaker coupling and stronger interference from multiple sources.

**Fig 11:**
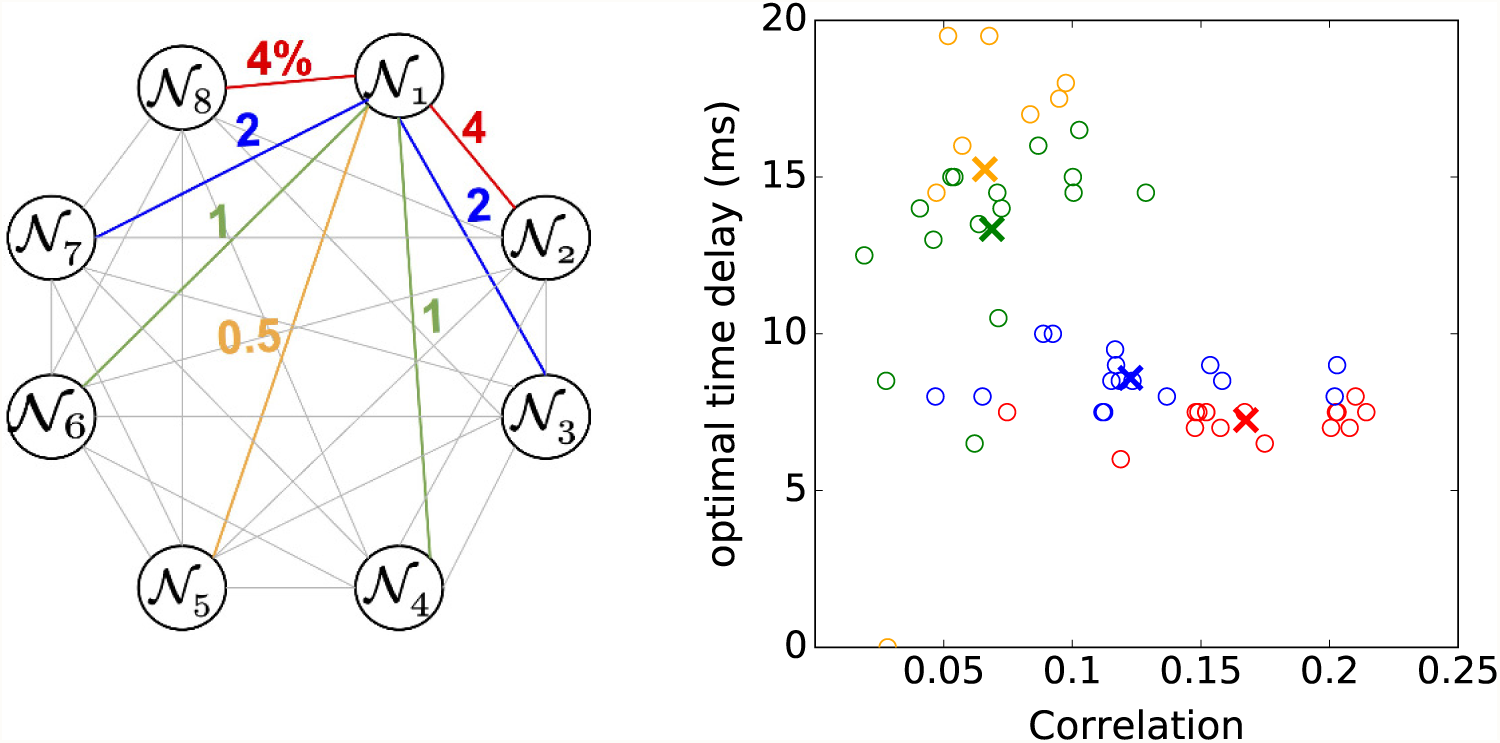
All-to-all coupled ring. Connectivities between two networks depend only on the distance between them; numbers on edges denote percentages of connectivity. The scatter plot shows the correlation and optimal time delays for all pairs of local populations (𝒩_*i*_,𝒩_*j*_), *i* ≠ *j*. Data are based off of one trial of 8 second simulation. The colors of the open circles indicate the distance between the pair: distance 1 in red, distance 2 in blue, distance 3 in green, and distance 4 in orange. For each specific pair, correlation is computed at the time delay that maximizes it. Crosses depict the average correlation and time delay for each distance group of pairs

Looking at the data points for groups separated by distances 1 and 2, we note that correlations had a significantly larger variance than optimal time delays, as was observed earlier in Figure 5 for a different question we were investigating.

## 5 Methods

### Spike synchrony index (SSI)

The following measures of synchrony were introduced in Chariker et al. (2018). We recall them for the convenience of the reader. Fix a window of length *w* ms. We used *w* = 4 which was optimal for studying gamma-band rhythms. For every excitatory spike that occurs at time *t*, we compute the fraction of the excitatory population that spiked within a window of *w* ms centered at *t*. These fractions are then averaged, giving the quantity we call SSI. Formally, if *N* is the size of the excitatory population and *F* = {*t*_*i*_, *i* = 1, 2, …} is the set of times during the simulation when an E neuron from the population spiked, then

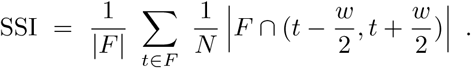

Here | · | denotes the cardinality of a set. If the system fires population spikes, most of which fall within a 2 ms interval, then SSI ∼ 1. Since most systems are nowhere close to being that synchronized, the following quotient gives more useful information:

Given a neuronal system, let SSI-null be the fraction of E-neurons spiking per *w*-sized bin if the system was firing perfectly homogeneously. That is, if *F* is the set of spike times during *L* seconds, then

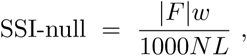

and the SSI-quotient is defined to be

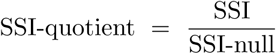

where the null system is assumed to have the same firing rate as the system in question. An SSI-quotient of 2, for example, means that compared to the corresponding null system, the system in question has, on average, twice as many spikes within a *w*-size window centered at each spike fired.

We stress that SSI-values are signatures of population activity. They measure the degree of synchrony of the population as a whole, without regard to which neurons fired which spikes or the level of correlated activity between specific pairs of neurons. SSI has similarities with spike field coherence (SFC), but differs because it considers spike-spike synchrony while SFC considers spike-local field potential synchrony (Chariker et al. 2018; Fries 2005; Chalk et al. 2010; Fries et al. 2001; Jia et al. 2013).

### Power spectral density

This is a widely used tool in the description of gamma-band activity in experiments. It is the Fourier transform of the spike density’s autocorrelation function (Wiener 1933). The spike density, *u*(*t*), is the the expected number of spikes/ms for an E neuron during the time bin [*t, t* + *Δt*). We compute the PSD using 4 seconds of simulation data and 200 ms overlapping windows as follows: let the population have *N* excitatory neurons. Consider a fixed time interval [0, *T*) which we divide into time bins, *B*_*n*_ = [(*n* − 1)*Δt, nΔt*) for *n* = 1, 2, … Then, the spike density is

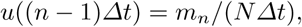

where *m*_*n*_ is the total number of excitatory spikes fired in *B*_*n*_. For ease, we refer to *u*((*n* − 1)*Δt*) as *u*(*n*). The discrete Fourier transform of *u*_*n*_ on (0, *T*) is given by the following:

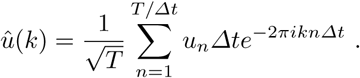

The PSD is then defined as |*û*(*k*)^2^|, and for each *k* represents the signal “power” concentrated at frequency *k*. For our computations, we used *T* = 0.2 s, *Δt* = 0.0025 s. The plots displayed in Figure 1e and the bottom of Figure 2a,b show averages of the PSD over sliding windows of length *T* in increments of 2.5 ms. In essence, the PSD was computed as described above for [0, *T*), [0.0025, *T* + 0.0025), … for the 4 seconds of simulation data. Then, the resulting PSDs were averaged (Chariker et al. 2018; Jenkins and Watts 1968).

## Discussion

This paper contains a theoretical study of correlations in spiking activity seen on the population level, without distinguishing between the behaviors of individual neurons. Using a network motif with multiple components, each one resembling a group of neurons connected by local circuitry, we studied how the spiking activity of two local populations are correlated on timescales of a few milliseconds.

### Take-home messages

The main points of this paper can be summarized as follows:

1. Spike-count correlations on the population level provide more robust metrics than similar correlations between pairs of neurons. They are an order of magnitude larger and have smaller variability. We offer the following explanation based on our analysis: Spike times of individual neurons are relatively random as can be seen from their ISI distributions. Collectively, neurons connected by local circuitry produce gamma rhythms when driven. Though the rhythm itself is generated from within the local population, we have found that the *timing* of spiking events is nontrivially influenced by external inputs, so that rhythms in target populations have a tendency to partially lock onto rhythms of source populations, producing the strong positive correlations seen.
2. In all of the situations considered, we have found that between source and target populations separated by no more than 2-3 synapses, there is a well defined notion of *response time*, i.e., there is a value of time delay, depending on network attributes but independent of initial condition, that maximizes the correlation. We showed that when subsampling, response times stabilize with as few as 10 neurons, unlike correlations, which are highly dependent on sample size.
3. We have found that synchrony in the source network produces higher correlations downstream, as does higher connectivity between populations. The situation *vis a vis* firing rates is less clear: contrary to what is commonly thought, we found that neither synchrony of the source network nor stronger connectivity necessarily produces higher firing in the target network.
4. We have also investigated correlations in some network motifs that occur in the real brain. Our main findings are (i) correlations between source and target decay rapidly with the number of intermediate layers; and (ii) in the presence of feedback or multiple pathways, incompatible response times may lead to lowered correlations, even as recurrent excitation and/or connectivity are increased.

### Gamma-band rhythms vs regular oscillations

As noted in our first take-home message, the alignment of gamma-band rhythms contributes significantly to the positive correlations between local populations. A distinguishing feature of this work is that we have used models that produce relatively realistic gamma-band rhythms (Sect. 1). In the real brain, gamma rhythms are broad-band, episodic, varying in frequency and in phase (Henrie and Shapley 2005; Xing et al. 2012). They are very far from regular oscillatory behavior, and produce correlations with a distinctive character.

For example, two populations that are oscillating periodically can lock onto one another perfectly when their frequencies coincide, and if their frequencies are incommensurate, then there will necessarily be substantial periods of incoherence. With realistic gamma properties, the situation is more nuanced: Gamma rhythms of the target network can partially lock onto that of the source network irrespective of the peak frequencies of the two populations (Sect. 3.1). This alignment, however, is not maintained, because realistic gamma rhythms degrade and resynchronize. This enables a population to be correlated to two independent sources (Sect. 4, Motif B), while purely oscillatory dynamics can never lock on to two other oscillations that differ either in frequency or in phase. The same degradation causes correlations to decay with distance from the source (Sect. 4, Motif A), unlike purely oscillatory systems, which can pass a signal perfectly along an arbitrarily long feedforward chain.

### Related Works

There is other literature on correlations between brain regions, not surprisingly as these are hypothesized to be important quantities for information transfer. Some related experimental work was mentioned in the Introduction. We postponed the discussion of theoretical results until now so that we can discuss their relation to our work.

A very influential idea proposed is *communication through coherence* (Fries 2005, 2015). We do not know whether by “communication” the author of Fries (2005, 2015) was referring to higher firing rates or higher correlations. We sought to clarify this in our study; the results are reported in Item 3 of the take-home messages.

Two theoretical papers with significant overlap with ours are Battaglia et al. (2012) and ter Wal and Tiesinga (2017). Similar to the present paper, Battaglia et al. (2012) investigated correlations between components in multi-component networks, but their goals were different, and they mostly considered averaged potentials (as opposed to the actual spikes) and rate models. See also Sancristóbal et al. (2014). ter Wal and Tiesinga (2017) focused on phases, including phase differences between stimuli and networks, as well as between source and target networks. Spike events in their networks (modeled on PING) seemed fairly periodic, facilitating the study of phases. The idea of phases appeared also in our work (Sect. 4, Motif C), though phases are less well defined because of the more episodic nature of our gamma rhythms. The more nuanced properties of realistic gamma and their roles in correlations between brain regions are a novel feature of our paper not considered in Battaglia et al. (2012) nor ter Wal and Tiesinga (2017), nor in any other paper that we know of.

Also related are papers that considered oscillatory feedforward input signals to populations, i.e. correlations between a population’s response and a signal, e.g. Gielen et al. (2010); Börgers and Kopell (2008). There are also studies that modeled the target population as signal filters, e.g. Akam and Kullmann (2012).

Further afield are papers that modeled the entrainment of single or pairs of neurons, e.g. Gielen et al. (2010); we have focused on population activity though subsampling was also discussed. Finally, we mention that there is a literature on correlations within single populations, as in (Brunel 2000; Brunel and Wang 2003; Rosenbaum and Doiron 2014). These papers have no overlap with us as they do not discuss correlations between distinct local populations.

### Implications and potential applications

Functions of gamma-band activity in the transmission of information or in the organization of brain activity have been hypothesized (Pesaran et al. 2002; Buzsáki and Wang 2012; Gray 1999; Fries 2015). If these hypotheses are valid, then correlations measure the effectiveness in the transmission of gamma patterns, which affects the above activities. It has also been documented that diseases and drugs lead to altered gamma characteristics. For example, individuals with disorders such as schizophrenia are known to exhibit abnormal gamma-band activity (Gonzalez-Burgos et al. 2010, 2015; Uhlhaas and Singer 2010), and drugs such as anesthesia are known to produce different gamma patterns (McCarthy et al. 2012). See also Chariker et al. (2018). Analysis along the lines carried out in this paper may shed light on whether (and if so, how) abnormal gamma rhythms will manifest themselves in unusual correlation values between local populations. Conversely, unusual correlation values may be indicative of abnormal cortical states.

Lastly, the population metrics proposed in this paper (with different window sizes) are applicable to oscillations in the beta, alpha, and theta band. It has been shown that these rhythms also lead to correlations measured on different time scales (Burke et al. 2013). Unlike gamma rhythms, which are largely (though not completely) produced in local populations, it has been proposed that these slower rhythms are more likely top-down. The ideas of this paper, namely correlations and response times, are equally relevant for such processes as they offer measures of how top-down effects are propagated.

## Acknowledgements

We would like to thank Wolf Singer for valuable discussions, and Robert Shapley for providing helpful comments on the manuscript.

## Supplementary Information

### Model equations and specifications

To connect our model neurons, we use the following rules: Each *E*-neuron is postsynaptic to *ζ*_1_ *E*-neurons where *ζ*_1_ is a Gaussian random variable with mean *μ* = 80 and standard deviation *σ* = 15 truncated at one SD. It is also postsynaptic to *ζ*_2_ *I*-neurons where *ζ*_2_ has the same form as *ζ*_1_ but with *μ* = 50 and *σ* = 7.5. Each inhibitory neuron is post-synaptic to *ζ*_3_ *E*-neurons with *μ* = 240 and *σ* = 37.5, and to *ζ*_4_ *I*-neurons with *μ* = 50 and *σ* = 7.5. The random variables *ζ*_*i*_ are independent from each other and independent from neuron to neuron, and the sets of presynaptic neurons are chosen randomly.

The dynamics of individual neurons are determined by the following leaky integrate-and-fire (LIF) equations: The normalized membrane potential *V* of a neuron *n* is governed by

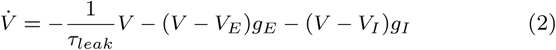

where time is measured in ms, and *V* is normalized so that when it reaches 1, the neuron *n* fires an action potential following which its membrane potential is immediately reset to 0. Eq. (2) has three constants: *τ*_*leak*_ = 20 ms is the leak rate, and 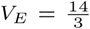 and 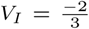 are excitatory and inhibitory reversal potentials in normalized units. These are accepted biophysical constants (Koch 1999; Chariker et al. 2016). On the right side of Eq. (2) are two functions *g*_*E*_ (*t*) and *g*_*I*_ (*t*), the excitatory and inhibitory conductances of neuron *n*, the dynamics of which are described below. When an action potential is fired, we assume a refractory period of 2 ms during which the potential does not update – though the conductances *g*_*E*_ and *g*_*I*_ continue to evolve.

The equations governing the evolution of *g*_*E*_ and *g*_*I*_ are as follows: We assume the neuron *n* in question is of type *Q* ∈ {*E, I*}, and let *M*_*E*_ (*n*) and *M*_*I*_ (*n*) be its sets of presynaptic excitatory and inhibitory neurons respectively. Then

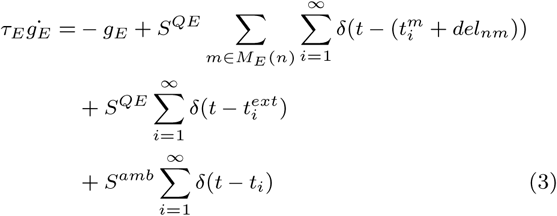

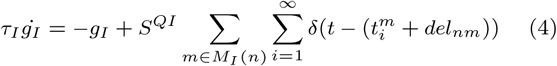

The E- and I-conductances *g*_*E*_ and *g*_*I*_ are elevated when neuron *n* receives some excitatory, respectively inhibitory, input, and *τ*_*E*_ and *τ*_*I*_ are the rates at which they decay to zero. The excitatory input to a neuron comes from three sources: synaptic input from other neurons within the local population, synaptic input from external sources such as other regions of cortex, and an “ambient” source representing unmodeled sources of neurotransmitters. The inhibitory input to a neuron comes from only one source, the synaptic input from other neurons in the population.

The term with a double-sum on the right side of Eq. (3) represents the synaptic excitatory input received by neuron *n*: We assume that the coupling weight between neurons depends only on their types, i.e. E or I, and *S*^*QE*^ is the coupling weight from E-neurons to neurons of type *Q*. The sequence 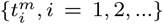 are the times when neuron *n* successfully receives synaptic input from neuron *m*. We assume a synaptic failure rate of 50%, which means that a neuron will have a 50% chance of successfully receiving a spike when a presynaptic neuron fires an action potential. We also assume that once a spike is received, the change in conductance is not instantaneous, i.e. there is some delay between presynaptic firing and postsynaptic response. This is modeled with the *del*_*nm*_ terms. In case there is any notational ambiguity, our use of the delta function is intended to say that if a spike from neuron *m* ∈ *M*_*E*_ (*n*) is received at time 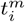 to neuron *n*, then *g*_*E*_ will increase by the amount 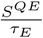at time 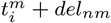. Eq. (4) is to be interpreted analogously.

Both excitatory and inhibitory neurons also receive external excitatory drive in the form of Poisson spikes arriving at random times 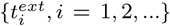, which are independent from neuron to neuron. The drive rates, *λ*_*E*_ and *λ*_*I*_ for excitatory and inhibitory receiving neurons, respectively, are variables we will modify and observe. Each time such a Poisson kick is received, the excitatory conductance of a neuron of type *Q* increases by 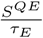.

Lastly, each neuron receives an ambient drive, in the form of Poisson kicks arriving at the times {*t*_*i*_, *i* = 1, 2, …}. This is meant to model resting state drive from neurotransmitters such as acetylcholine, and is represented by the term 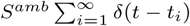 in Eq (3). The change in conductance caused by these kicks has magnitude *S*^*amb*^, different from *S*^*QE*^.

### Parameters used

We have tried to use parameters that are both consistent with what is accepted biologically and that produce the expected firing regimes in background and driven cases. Starting from the parameters in Chariker and Young (2015); Chariker et al. (2016), we made adjustments to accommodate for the smaller size of the present model network. The simulations used in this paper were produced using *S*^*EE*^ = .0255, *S*^*IE*^ = .0095, *S*^*EI*^ = .054, and *S*^*II*^ = .0275. We set the excitatory and inhibitory decay times to *τ*_*E*_ = 2ms and *τ*_*I*_ = 4ms, respectively. As to the values for *del*_*nm*_, if *n* or *m* is an inhibitory neuron, we set *delnm* = 1 ms. If both *n* and *m* are excitatory, we assume *delnm* is uniformly distributed on [1, 2.3] ms, to reflect the fact that exictatory to excitatory synapses range from axodendritic to axosomatic. The delays are set by pairs of neurons, and remain constant throughout the simulation.

The ambient drive rate is *λ*_*amb*_ = .433 kicks per ms, and *S*^*amb*^ is set to .003. The ambient drive was chosen such that, on average, around 10 percent of the total amount of excitatory input to each neuron was ambient.

Finally, the external Poisson drive rates *λ*_*E*_ and *λ*_*I*_ were allowed to vary to produce the desired firing rates in background and driven regimes. To simulate background conditions, we set 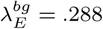 spikes per ms and 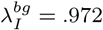 spikes per ms (consistent with the numbers of presynaptic E-neurons for E and I cells from within the local population). This gives an average spiking rate of about 2.5 spikes per second for excitatory neurons and 7.5 spikes per second for inhibitory neurons. We vary the external drive rates, *λ*_*E*_ and *λ*_*I*_ by choosing a constant, *c* ≥ 1, and setting 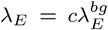and 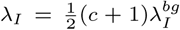. The factor of increase in *λ*_*E*_ and *λ*_*I*_ are different to best preserve the firing rate ratio of inhibitory neurons spiking 3-4 more times per second than excitatory neurons spike. We think of *c* = 1 as background and *c* = 2.25 as strongly driven, as they elicit firing rates consistent with the range seen in, for example, the visual cortex.

### Connecting two local populations

(details for the construction in Sect. 2.1) To model one neuronal layer feeding into another, we first create two separate networks, 𝒩_1_ and 𝒩_2_, to be thought of as the “source” and “target” networks respectively, following the specifications in section 1. For *m* = 1, 2, we let *εm* and ℐ*m* be the set of excitatory and inhibitory neurons in the network 𝒩*m*. We then replace a certain percentage of the external drive to neurons in *ε*_2_ and ℐ_2_ by excitatory input from “feedforward” connections from *ε*_1_.

More precisely, suppose we wish to have a fraction *p* ∈ (0, 1) of the total excitatory drive to neurons in 𝒩_2_ be feedforward drive from 𝒩_1_. In the text, we refer to this number *p* as the *connectivity from* 𝒩_1_ *to* 𝒩_2_. The construction is carried out in four steps:

1. First we compute for E-neurons in 𝒩_2_ the mean number, *f*_*E*_, of presynaptic neurons from *ε*_1_. Let the external drive rates to E-neurons in 𝒩*m* be *λE,m* and their firing rates be *r*_*E,m*_ for *m* = 1, 2. We solve for *f*_*E*_ in

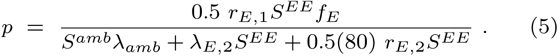 The denominator represents the total excitatory “drive” received by an E-neuron in 𝒩_2_, counting the “drive” from each of the three sources as the product of the number of spikes and the synaptic weight of each spike. In the last term, for example, 0.5 is the synaptic failure rate, 80 is the mean number of presynaptic E-neurons from within the local population, and *r*_*E,*2_ is the mean firing rate of the E-population in 𝒩_2_. The numerator in Eq. (5) is the total “drive” provided to the neuron in question by the E-population from 𝒩_1_. The mean number of presynaptic E-cells from 𝒩_1_ for each I-cell in 𝒩_2_, denoted *f*_*I*_, is computed similarly. For the regime depicted in Sect. 1.1, the computations above yield *f*_*E*_ =17.3 and *f*_*I*_ = 47 at *p* = 0.1, to give an example.
2. Next we set the connections between 𝒩_1_ and 𝒩_2_: Let *ζ* be the random variable in the “Model equations and specifications” section above. For E-neurons in 𝒩_2_, we set *μ* = *f*_*E*_ and *σ* = 30*p* (so that at *p* = 0.1, *μ* = 17.3 and *σ* = 3, a variance roughly commensurate with those within local populations). For I-neurons in 𝒩_2_, *μ* = *f*_*I*_ and *σ* = 90*p*. The random variable *ζ* is redrawn for each neuron in 𝒩_2_, and the neuron is then postsynaptic to *ζ* randomly selected neurons from *ε*_1_.
3. Having added the feedforward drive to 𝒩_2_, we now need to remove the corresponding fraction of *λ*_*E,*2_ and *λI,*2, in such a way that the total excitatory current from external sources received by neurons in 𝒩_2_ is the same as it would have been in the absence of the “feedforward” drive from 𝒩_1_. This completes the network architecture for the source-target pair 𝒩_1_ → 𝒩_2_.
4. . Finally, spikes fired by neurons in *ε*_1_ are received by E- and I-neurons in 𝒩_2_ the same way that excitatory spikes from within 𝒩_2_ are received — except for an additional transmission time of *δ*^+^ ms. This parameter is defined in the the text, within the context of different network motifs.

## Notes

This research is partially supported by National Science Foundation Grant DMS-1363161

## References

Akam, T. E. and Kullmann, D. M. (2012). Efficient “communication through coherence” requires oscillations structured to minimize interference between signals. PLOS Computational Biology, 8.

Baddeley, R., Abbott, L. F., Booth, M. C. A., Sengpiel, F., Freeman, T., Wakeman, E. A., and Rolls, E. T. (1997). Responses of neurons in primary and inferior temporal visual cortices to natural scenes. Proceedings of the Royal Society of London B, pages 1775–1783.

Bastos, A. M., Vezoli, J., and Fries, P. (2015). Communication through coherence with inter-areal delays. Current Opinion in Neurobiology, 31:173–180.

Battaglia, D., Witt, A., Wolf, F., and Geisel, T. (2012). Dynamic effective connectivity of inter-areal brain circuits. PLOS Computational Biology, 8.

Binzegger, T., Douglas, R. J., and Martin, K. (2009). Topology and dynamics of the canonical circuit of cat v1. Neural Networks, 22:1071–1078.

Börgers, C. and Kopell, N. J. (2008). Gamma oscillations and stimulus selection. Neural Computation, 20:383–414.

Bosman, C. A., Schoffelen, J.-M., Brunet, N., Oostenveld, R., Bastos, A. M., Womelsdorf, T., Rubehn, B., Stieglitz, T., Weerd, P. D., and Fries, P. (2012). Attentional stimulus selection through selective synchronization between monkey visual areas. Neuron, 75:875–888.

Brunel, N. (2000). Dynamics of sparsely connected networks of excitatory and inhibitory spiking neurons. Journal of Computational Neuroscience, 8:183–208.

Brunel, N. and Wang, X.-J. (2003). What determines the frequency of fast network oscillations with irregular neural discharges? i. synaptic dynamics and excitation-inhibition balance. Journal of Neurophysiology, 90:415–430.

Burke, J. F., Zaghloul, K. A., Jacobs, J., Williams, R. B., Sperling, M. R., Sharan, A. D., and Kahana, M. J. (2013). Synchronous and asynchronous theta and gamma activity during episodic memory formation. Journal of Neuroscience, 33:292–304.

Buzsáki, G. (2011). Rhythms of the brain. Oxford University Press.

Buzsáki, G. and Wang, X.-J. (2012). Mechanisms of gamma oscillations. Annu Rev Neurosci., 35:203–225.

Chalk, M., Herrero, J. L., Gieselmann, M. A., Delicato, L. S., Gotthardt, S., and Thiele, A. (2010). Attention reduces stimulus-driven gamma frequency oscillations and spike field coherence in v1. Neuron, 66:114–125.

Chariker, L., Shapley, R., and Young, L.-S. (2016). Orientation selectivity from very sparse lgn inputs in a comprehensive model of macaque v1 cortex. The Journal of Neuroscience, 36(49):12368–12384.

Chariker, L., Shapley, R., and Young, L.-S. (2018). Rhythm and synchrony in a cortical network model. The Journal of Neuroscience, 38(40):8621–8634.

Chariker, L. and Young, L.-S. (2015). Emergent spike patterns in neuronal populations. Journal of Computational Neuroscience, 38:203–220.

Diesmann, M., Gewaltig, M.-O., and Aertsen, A. (1999). Stable propagation of synchronous spiking in cortical neural networks. Nature, 402:529–533.

Fries, P. (2005). A mechanism for cognitive dynamics: neuronal communication through neuronal coherence. TRENDS in Cognitive Sciences, 9:10496–10504.

Fries, P. (2015). Rhythms for cognition: Communication through coherence. Neuron, 88:220–235.

Fries, P., Reynolds, J. H., Rorie, A. E., and Desimone, R. (2001). Modulation of oscillatory neuronal synchronization by selective visual attentive. Science, 291:1560–1563.

Gielen, S., Krupa, M., and Zeitler, M. (2010). Gamma oscillations as a mechanism for selective information transmission. Biological Cyber-netics, 103:151–165.

Gonzalez-Burgos, G., Cho, R. Y., and Lewis, D. A. (2015). Alterations in cortical network oscillations and parvalbumin neurons in schizophrenia. Biological Psychiatry, 77:1031–1040.

Gonzalez-Burgos, G., Hashimoto, T., and Lewis, D. A. (2010). Alterations of cortical gaba neurons and network oscillations in schizophrenia. Current Psychiatry Reports, 12:335–344.

Gray, C. M. (1999). The temporal correlation hypothesis of visual feature integration: still alive and well. Neuron, 24:31–47.

Gray, C. M., König, P., Engel, A. K., and Singer, W. (1989). Oscillatory responses in cat visual cortex exhibit inter-columnar synchronization which reflects global stimulus properties. Nature, 338:334–337.

Gray, C. M. and Singer, W. (1989). Stimulus-specific neuronal oscillations in orientation columns of cat visual cortex. Proceedings of the National Academy of Sciences of the United States of America, 86:1698–1702.

Grinvald, A., Lieke, E. E., Frostig, R. D., and Hildesheim, R. (1994). Cortical point-spread function and long-range lateral interactions revealed by real-time optical imaging of macaque monkey primary visual cortex. Journal of Neuroscience, 14:2545–2568.

Henrie, J. A. and Shapley, R. (2005). LFP power spectra in V1 cortex: The graded effect of stimulus contrast. Journal of Neurophysiology, 94:479–490.

Jenkins, G. and Watts, D. G. (1968). Spectral analysis and its applications. Holden Day.

Jia, X., Smith, M. A., and Kohn, A. (2011). Stimulus selectivity and spatial coherence of gamma components of the local field potential. The Journal of Neuroscience, 31:9390–9403.

Jia, X., Tanabe, S., and Kohn, A. (2013). Gamma and the coordination of spiking activity in early visual cortex. Neuron, 77:762–774.

Khawaja, F. A., Tsui, J. M., and Pack, C. C. (2009). Pattern motion selectivity of spiking outputs and local field potentials in macaque visual cortex. Journal of Neuroscience, 29:13702–13709.

Koch, C. (1999). Biophysics of Computation: Information Processing in Single Neurons. Oxford University Press.

Kuhn, A., Aertsen, A., and Rotter, S. (2003). Higher-order statistics of input ensembles and the response of simple model neurons. Neural Computation, 15:67–101.

McCarthy, M. N., Ching, S. N., Whittington, M. A., and Kopell, N. (2012). Dynamical changes in neurological diseases and anesthesia. Current Opinion in Neurobiology, 22:693–703.

Ostojic, S. (2011). Interspike interval distributions of spiking neurons driven by fluctuating inputs. Journal of Neurophysiology, pages 361–373.

Pesaran, B., Pezaris, J. S., Sahani, M., Mitra, P. P., and Andersen, R. A. (2002). Temporal structure in neuronal activity during working memory in macaque parietal cortex. Nature Neuroscience, 5:805–811.

Pikovsky, A. (2001). Synchronization: A Universal Concept in Nonlinear Sciences. Cambridge University Press.

Rangan, A. V. and Young, L.-S. (2013). Emergent dynamics in a model of visual cortex. Journal of Computational Neuroscience, 35:155–167.

Rosenbaum, R. and Doiron, B. (2014). Balanced networks of spiking neurons with spatially dependent recurrent connections. Physical Review X, 4.

Salazar, R. F., Dotson, N. M., Bressler, S. L., and Gray, C. M. (2012). Content-specific fronto-parietal synchronization during visual working memory. Science, 338:1097–1100.

Sancristóbal, B., Vicente, R., and Garcia-Ojalvo, J. (2014). Role of frequency mismatch in neuronal communication through coherence. Journal of Computational Neuroscience, 37:193–208.

Sederberg, P., Kahana, M., Howard, M., Donner, E., and Madsen, J. (2003). Theta and gamma oscillations during encoding predict subsequent recall. Journal of Neuroscience, 23:10809–10814.

Sincich, L. C. and Horton, J. C. (2005). The circuitry of V1 and V2: integration of color, form, and motion. Annual Review of Neuroscience, 28:303–326.

Ter Wal, M. and Tiesinga, P. H. (2017). Phase difference between model cortical areas determines level of information transfer. Frontiers in Computational Neuroscience, 11.

Uhlhaas, P. J. and Singer, W. (2010). Abnormal neural oscillations and synchrony in schizophrenia. Nature Reviews Neuroscience, 11:100–113.

Whittington, M. A., Traub, R. D., Kopell, N., Ermentrout, B., and Buhl, E. H. (2000). Inhibition-based rhythms: experimental and mathematical observations on network dynamics. International Journal of Psychophysiology, 38:315–336.

Wiener, N. (1933). The Fourier integral and certain of its applications. Cambridge University Press.

Xing, D., Shen, Y., Burns, S., Yeh, C. I., Shapley, R., and Li, W. (2012). Stochastic generation of gamma-band activity in primary visual cortex of awake and anesthetized monkeys. Journal of Neuroscience, 32:13873–13880.

Zandvakili, A. and Kohn, A. (2015). Coordinated neuronal activity enhances corticocortical communication. Neuron, 87:827–839.

